# The extent and impact of variation in ADME genes in sub-Saharan African populations

**DOI:** 10.1101/2020.06.14.108217

**Authors:** Jorge da Rocha, Houcemeddine Othman, Gerrit Botha, Laura Cottino, David Twesigomwe, Samah Ahmed, Britt I. Drögemöller, Faisal M. Fadlelmola, Philip Machanick, Mamana Mbiyavanga, Sumir Panji, Galen E.B. Wright, Clement Adebamowo, Mogomotsi Matshaba, Michèle Ramsay, Gustave Simo, Martin C. Simuunza, Caroline T. Tiemessen, Sandra Baldwin, Mathias Chiano, Charles Cox, Annette S. Gross, Pamela Thomas, Francisco-Javier Gamo, Scott Hazelhurst, as members of the H3Africa Consortium

## Abstract

Investigating variation in genes involved in the *absorption, distribution, metabolism*, and *excretion* (ADME) of drugs are key to characterising pharmacogenomic (PGx) relationships. ADME gene variation is relatively well characterised in European and Asian populations, but African populations are under-studied – which has implications for safe and effective drug use in Africa.

The genetic diversity of ADME genes across sub-Saharan African populations is large. The Southern African population cluster is most distinct from that of far West Africa. PGx strategies based on European variants will be of limited use in African populations.

Although established variants are important, PGx must take into account the full range of African variation. This work urges further characterisation of variants in African populations including *in vitro* and *in silico* studies, and to consider the unique African ADME landscape when developing precision medicine guidelines and tools for African populations.

**Author summary:** The ADME genes are a group of genes that play a key role in absorption, distribution, metabolism and excretion of drugs. Variations in these genes can have a significant impact on drug safety and efficacy. Africa has a high level of genetic variation and is under-studied in drug development, which makes study of variations in these genes in African populations very important. Using a new data set of 458 high-coverage genomes from across Africa, we characterise the extent and impact of variation in the ADME genes, looking at both single nucleotide and copy number variations. We identified 343,368 variants, including 40,692 novel variants, and 930 coding variants which are predicted to have high impact on function. Our discovery curves indicate that there will be considerable value in sequencing more African genomes. Moreover, relatively few of these novel variants are captured on common genotyping arrays. We show that there is considerable diversity within Africa in some important genes, and this will have significant consequences for the emerging field of precision medicine in Africa.

## 1 Introduction and background

Pharmacogenomics (PGx) aims to improve drug safety and efficacy using genomic knowledge for genes involved in drug action [1] with a focus on genes that have important roles in drug safety, pharmacokinetics and pharmacodynamics. Genes involved in pharmacokinetics are typically defined by the role they play in the absorption, distribution, metabolism and excretion (ADME) of drug molecules.

Variation in ADME genes play an important role in determining the response to drug treatment in an individual patient. We characterise the extent and impact of variation in these genes in a novel, high-coverage whole genome sequence dataset from a diverse group of Africans.

ADME genes have different functions: (1) phase I metabolising enzymes, (2) phase II metabolising enzymes, (3) drug transporters and (4) modifiers. PharmaADME (http://pharmaadme.org) classifies the ADME genes in two classes. The 32 *core* genes have known biomarkers linked to ADME outcomes. For the 267 *extended* ADME genes, there is weaker evidence of functional consequences *in vitro* or *in vivo*, or they are important for a limited number of drugs only.

### Rationale

Currently the majority of patients studied in drug development programmes are of European or Asian ancestry. The African continent is the cradle of human origin and African populations are characterised by high genetic diversity and complex population structure. Despite this genetic variation, drug efficacy and safety have not been comprehensively studied in the populations of Sub-Saharan Africa (SSA) [2]. This is of specific relevance to SSA, where high burdens of disease are amplified by non-optimal treatment outcomes.

The particular diversity of ADME genes in SSA has been reported in some studies. Hovelson *et al*. [3] and Lakiotaki *et al*. [4] found that the greatest levels of coding ADME variation per personal haplotype were shown in some African populations sampled in the 1000 Genomes Project (KGP) data. Examples of the impact of this variation can be seen in *CYP2B6* and *CYP2D6* variation affecting efavirenz and primaquine respectively. An efavirenz dosage reduction has been recommended for HIV patients in SSA due to the high frequency of functional variants in the *CYP2B6* gene that result in a higher risk of adverse drug reactions [5]. Potential polymorphisms in the human cytochrome *CYP2D6* gene may negatively influence efficacy of primaquine, and significantly affect malaria elimination strategies [6, 7]. African specific variation in several genes may impact the PK of rosuvastatin, a drug used to treat hypercholesterolemia [8]. While these studies represent only a fraction of the continent, they serve to highlight the importance of future studies which are aimed at providing a more comprehensive overview of the landscape of ADME variation across Africa.

Therefore it is important to gain a better understanding of the variation that exists in ADME genes, both within and between different SSA populations. This information could be used to inform recommended drug dosage regimens for patients in SSA based on potential pharmacokinetic effects and consequently efficacy and safety. To date, no studies have systematically investigated ADME variation within a diverse set of African populations. We therefore aim to provide valuable information regarding the variation that exists in ADME genes, both within and between different SSA populations. This information could provide insight into drug efficacy and safety for patients in SSA and play a role in ensuring safe and efficacious treatments for the high burden of diseases in populations in SSA. We have characterised the extent and impact of variation in ADME genes in a novel, high-coverage whole genome sequence dataset from a diverse group of Africans.

## 2 Results

### 2.1 Description of Samples

Four hundred fifty eight high coverage whole genome sequences were used in the study as the primary data set (we call this the high coverage African ADME Dataset – HAAD). The foundation of this set were sequences generated by the Human Health and Heredity in Africa (H3A) consortium [9, 10]. The sources and countries of origins of the samples can be found in Table 1 and Fig 1. The population structure of participants in this study is broadly representative of speakers of Niger-Congo languages from West through South Africa. Representation from Nilo-Saharan and Afro-Asiatic populations is sparse. There also are few individuals of Khoe and San heritage, although significant admixture from Khoe and San speakers is found in Bantu-speakers in Southern Africa [11].

**Table 1.** Sources of high-coverage data sets used to form HAAD: 272 genomes were generated by a supplementary grant from the NIH to the H3A Consortium [9] for the primary purpose of designing a custom genotyping array; 140 were shared by African collaborators; and the rest are publicly available, 15 genomes came from the South African Human Genome Programme, and 31 genomes were from the Simons Genome Diversity Project.

| H3A Consortium Data: High Coverage |  |  |
| --- | --- | --- |
| Benin | University of Montréal | 50 |
| Burkina Faso | AWI-Gen | 33 |
| Botswana | BHP | 47 |
| Cameroon | University of Dschang | 26 |
| Ghana | AWI-Gen | 26 |
| Nigeria | Institute of Human Virology | 49 |
| South Africa | AWI-Gen | 100 |
| Zambia | University of Zambia | 41 |
| African collaborators: High coverage |  |  |
| South Africa | SA Human Genome Programme | 15 |
| South Africa | Cell Biology Research Lab, NICD/Wits | 40 |
| Public data sets |  |  |
| Various | Simons Foundation | 31 |

**Fig 1.**
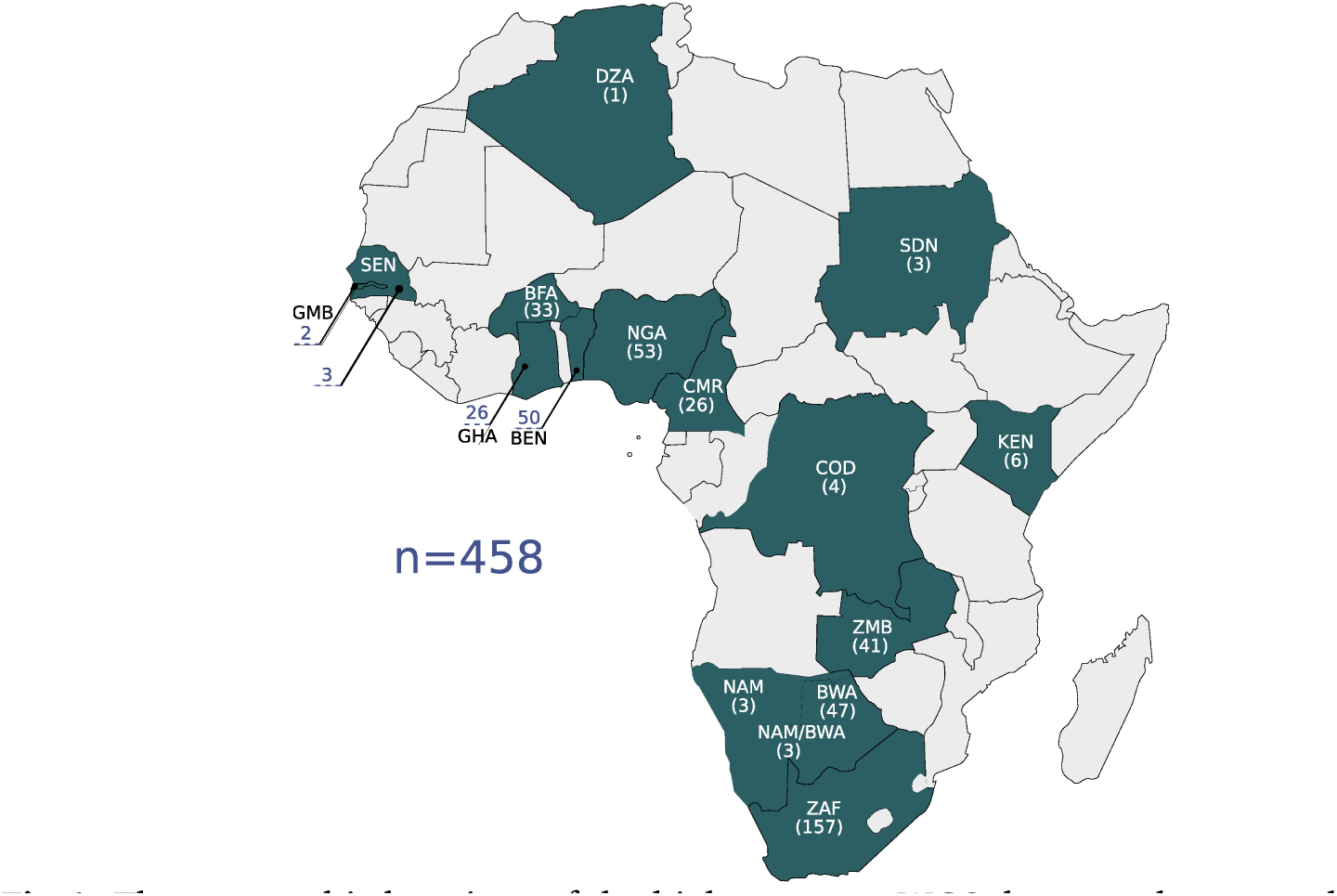
The geographic locations of the high coverage WGS data are shown on the map. Countries are referenced by their ISO 3166-1 alpha-3 code: BEN: Benin; BFA: Burkina Faso; BWA: Botswana; CMR: Cameroon; COD: Democratic Republic of the Congo; DZA: Algeria; GHA: Ghana; GMB: Gambia; KEN: Kenya; NAM: Namibia; NGA: Nigeria; SDN: Sudan; SEN: Senegal; ZAF: South Africa; ZMB: Zambia. The number of samples per country is shown in parentheses.

We supplement some analyses with African datasets from the KGP (we use KGA specifically to refer to the African genomes in KGP). As the KGP datasets are low coverage, not all analyses were performed with the KGA dataset in addition to HAAD.

### 2.2 Population structure

A principal component (PC) and structure analysis of our data shows high genome-scale variation and that we have significant breadth and depth of coverage of African genomic diversity across west, central and southern Africa, with lesser coverage in east Africa. The PC analysis of our data shows a strong correlation to geographical location (Fig 2 and supplementary section 1).

**Fig 2.**
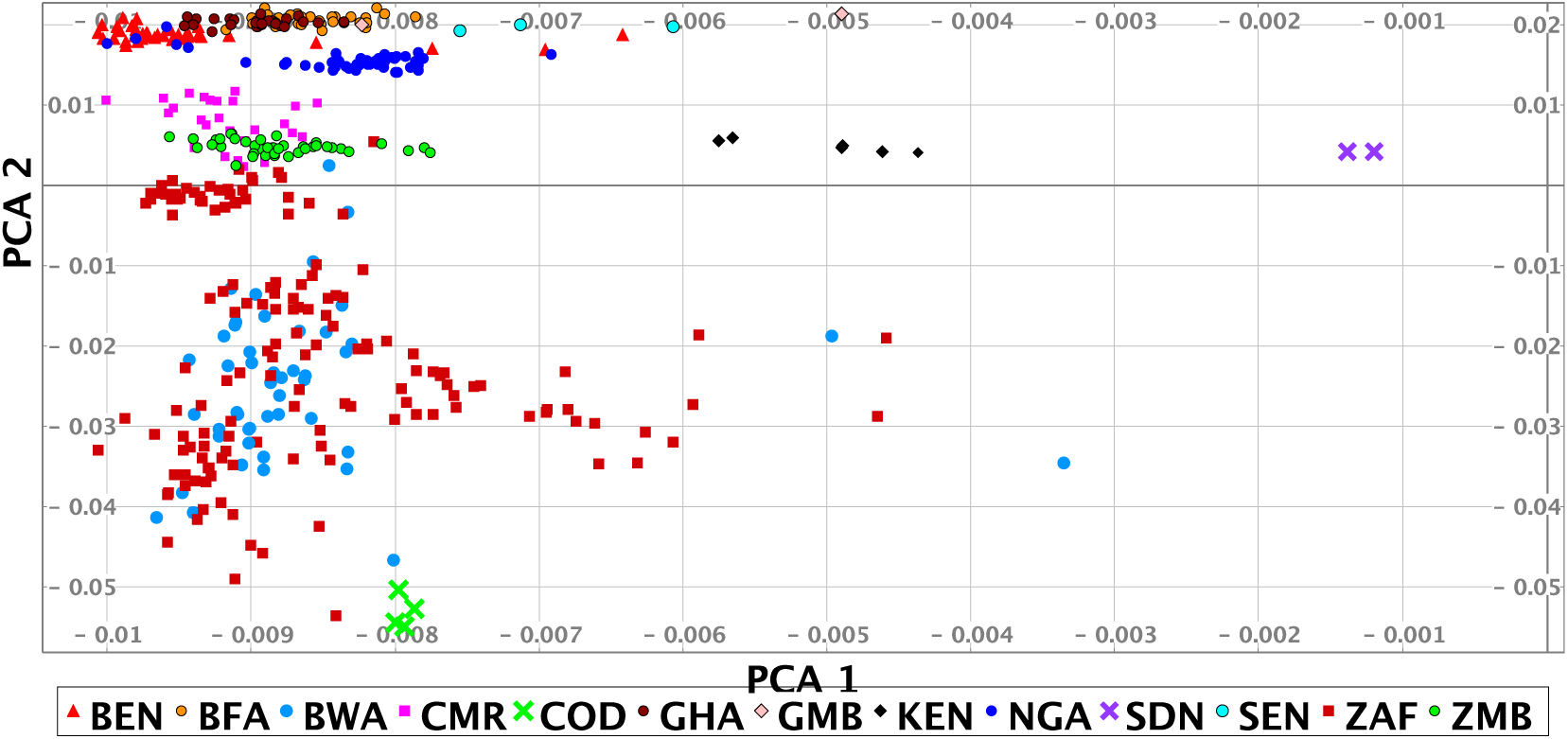
Principal component analysis of the HAAD (some outliers are omitted). Abbreviations for sources used are H3A (Human Health and Heredity in Africa Consortium), and SF (Simons Foundation Genome Diversity Project). The countries of origin and source of the samples shown in the PCA are: BEN/H3A, BFA/H3A, BWA/H3A+SF, CMR/H3A, COD/SF, GHA/H3A, GMB/SF, KEN/SF, NGA/H3A+SF, SEN/SF, ZAF/H3A+Tiemessen Lab+SF+South African Human Genome Programme, ZMB/H3A. Country codes given in Fig 1

To explore diversity between different African regions we clustered the studied population together with reference data sets using PC data (see *Methods*, Table 2). The PC analysis shows that the HAAD samples fall broadly into three groups: West (Ghana, Burkina Faso, Nigeria), South/Central (Cameroon, Zambia, Botswana, South Africa), South (Botswana, South Africa) African populations. The variability in the Southern group primarily arises through differential admixture between Bantu, Khoe and San speakers. There is a *Far West* group comprising individuals in HAAD and KGA from Gambia, Senegal and Sierra Leone. There are also a few individuals from other African regions. Note that there is significant diversity within countries; and in some cases overlap between countries – e.g. some participants that we label as “South/Central” live to the south of some participants in the “Southern” group.

**Table 2.** Clusters within Africa, including the number of individuals in each cluster. Clusters include both HAAD and 1000 Genomes African population data.

| Identifier | Number | Region |
| --- | --- | --- |
| SA | 166 | Southern Africa |
| SC | 172 | South/Central Africa |
| KS | 5 | Khoe and San |
| FW | 185 | Far West Africa |
| WE | 309 | West Africa |
| O | 5 | Outliers |

### 2.3 Overall Characterisation of ADME variation

Gene-based genetic variation for the core and extended ADME gene categories was assessed for composition and type, including introns, upstream and downstream flanking regions (Fig 3). Comparisons were made between the HAAD dataset and the KGA dataset, which represent samples in the joint called HAAD and African 1000 Genomes Project populations respectively (Methods 5.3.1). In ADME core genes, we counted a total of 40,714 and 36,088 variants for HAAD and KGA data respectively while for the extended ADME genes there were 274,798 and 243,022 variants respectively. Intronic variants are most common overall with about the same proportions in both HAAD and KGA datasets of 80% and 77% (for both core and extended genes) respectively. A significant number of variations appear in 3’ untranslated (3’ UTR) and 5’UTR regions. Coding region variants (non-synonymous and synonymous as annotated by VEP v92.0) do not overlap completely between HAAD and KGA groups. For core genes there were 423 coding variants common to both HAAD and KGA datasets, 288 coding variants unique to HAAD, and 252 unique to KGA. For extended genes, there were 17,148 coding variants common to HAAD and KGA, 2,850 unique to HAAD, and 2,318 unique to KGA respectively. Care should be taken in comparing HAAD and KGA data because of the different depth of sequencing.

**Fig 3.**
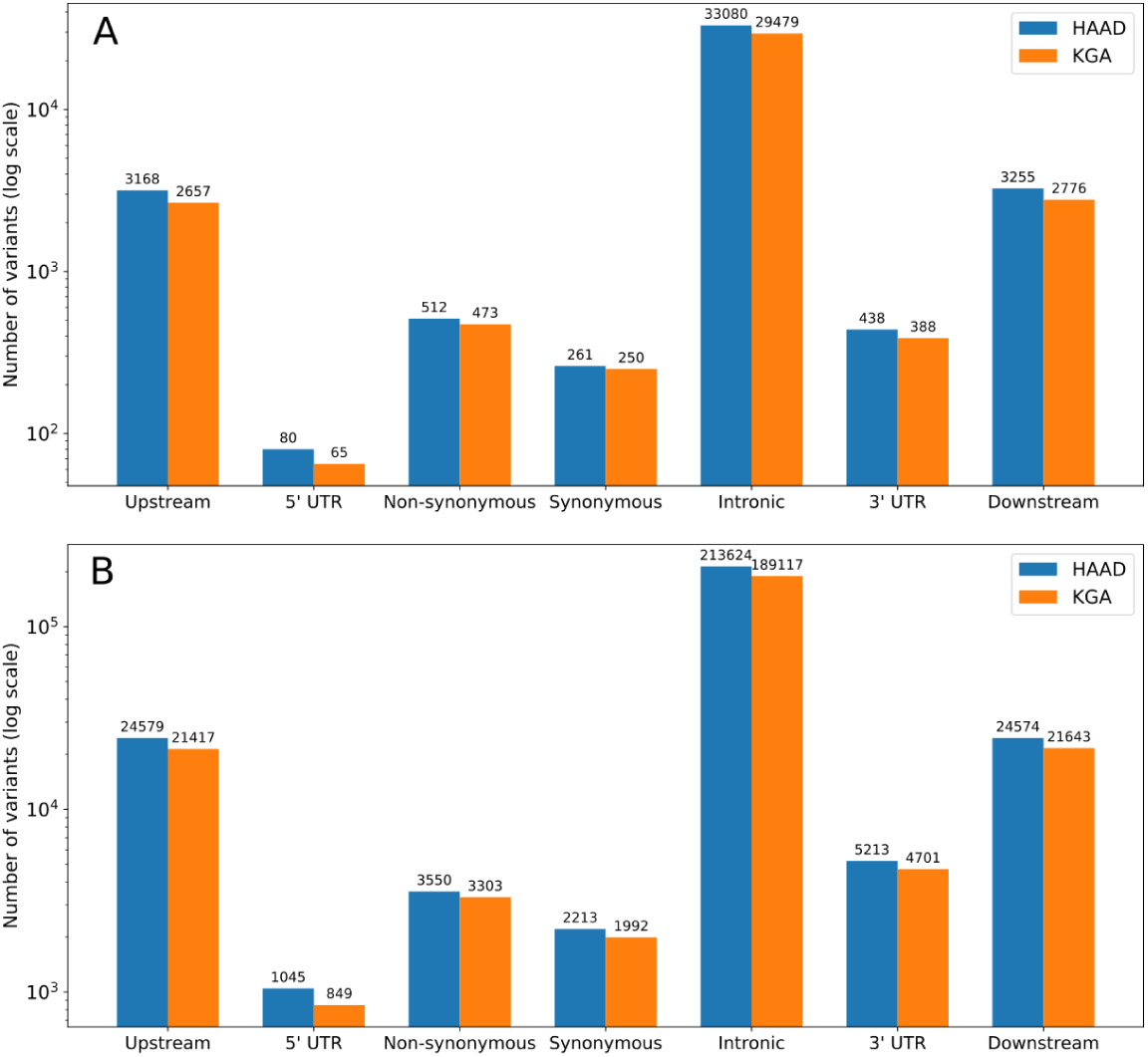
Distribution of variant types (as defined by SNPeff annotation) across core (A) and extended (B) ADME gene regions. HAAD (N=458) represents those samples in the jointly called set from the H3A Consortium data, Simons Foundation, SAHGP and Tiemessen Labs, and KGA (*n*=506) represents the African 1000 Genomes Project populations from the jointly called set. Upstream and downstream regions are represented by 10kb flanks from gene start and end respectively.

The importance of using and generating African datasets like ours can be seen in our discovery curves which show the increase in the number of variants found in the core ADME genes as more genomes are included in the study (the results for the extended genes are not shown but are similar). Fig 4 compares our data set to 1000 Genomes African and European populations. The diversity of African populations compared to European populations is clear and consistent with previous literature [3]. We believe that the increased richness of our data compared to 1000 Genomes African data is partially due to the fact that our data is high-coverage. This richness is also likely to be driven by the significant numbers of southern African genomes that have significant Khoe and San ancestry (see [11] for some discussion) as well some diverse samples from the Simons Foundation. Fig 5 shows the discovery curve for the combined African (HAAD and KGA) dataset. Although the curve has started to plateau, the results show that combining the data sets has value and sampling more Africans and more diverse African groups not yet properly captured will reveal considerably more variants.

**Fig 4.**
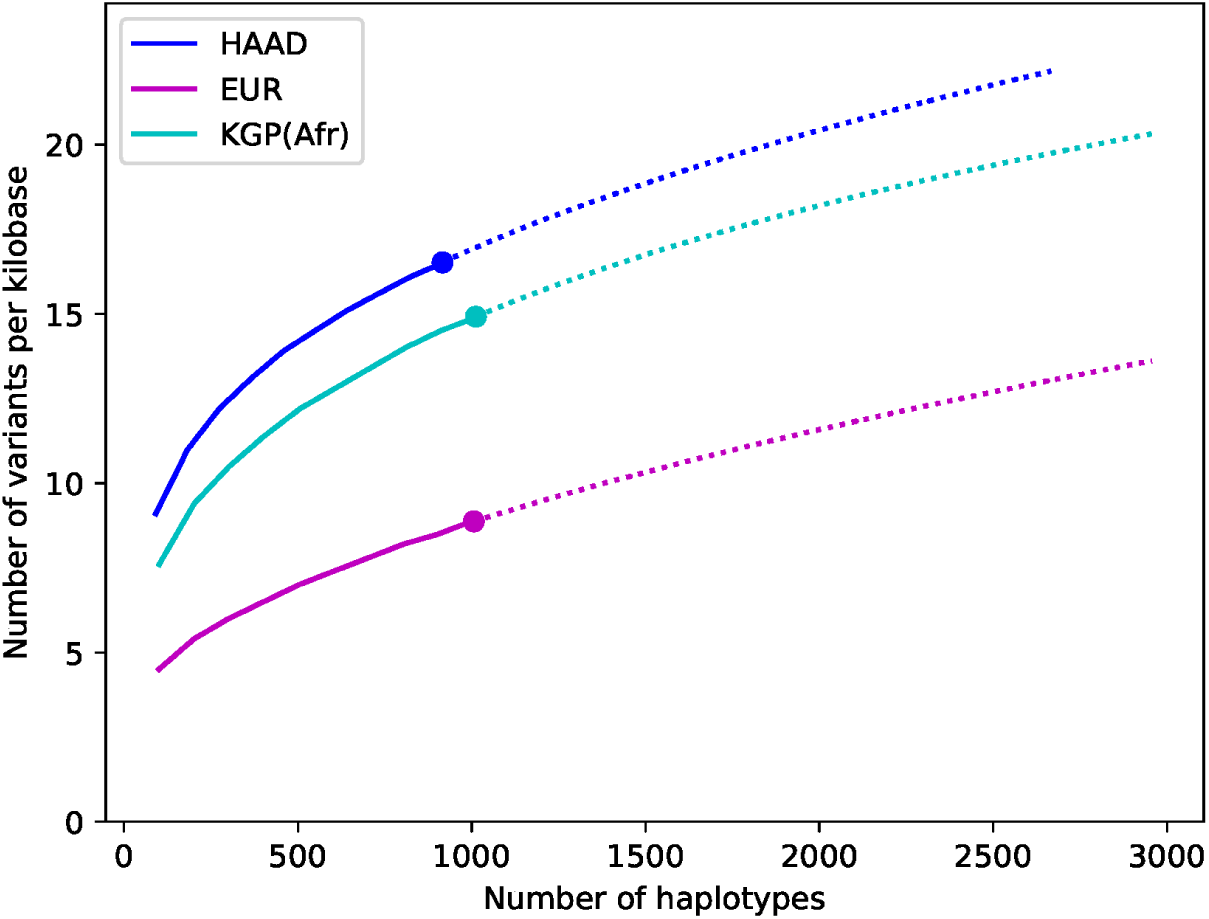
Comparative discovery curves of variants in the core ADME genes (including flanks) for the HAAD, and the 1000 Genomes Africa and European data. The results show for a given number of haplotypes the number of variants seen per kilobase. The actual results are shown as a large dot, sub-samples by a solid line, and projections by a dotted line. Sub-samples values are computed averaging over 50 different randomly sampled subsets for intermediate values. Projection is computed using a 3rd order jackknife projection [12].

**Fig 5.**
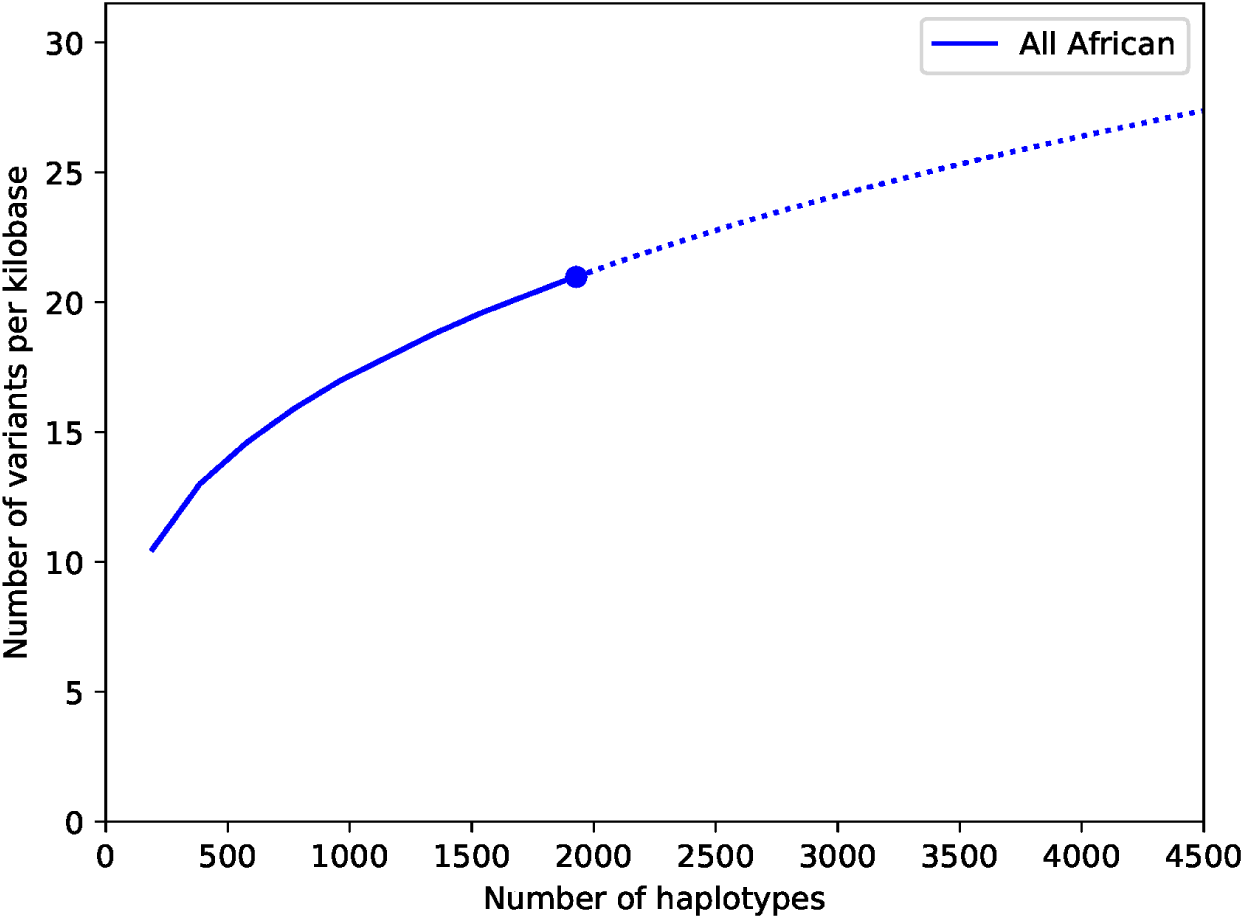
Discovery curve of variants in the core ADME genes in the combined HAAD and KGA datasets.

### 2.4 Annotation of high impact coding variants

To annotate ADME genes we used the output of an ADME gene optimised annotation schema. This schema uses five prediction tools, and variants meeting score cutoffs for all five are of the highest confidence for functional impact. We identified 930 high impact variants (HI-vars) for 247 ADME genes (from a total of 299 ADME genes) of which 29 are core genes and 218 are extended genes. Of the core genes, seven members of the cytochrome P450 (CYP450) family (*CYP1A1, CYP1A2, CYP2B6, CYP2C8, CYP2C19, CYP2A6, CYP2D6*) were among those with the highest count of high impact variants. Highest counts of the CYP450 genes were seen in *CYP1A1* and *CYP2D6* with 12 and 10

HI-vars of respectively. The ATP-Binding Cassette (ABC) transporter gene, *ABCB5* showed the highest number of HI-vars overall numbering 20. We also counted three members of ABCC transporter family and three other members of the CYP450 family in the 10 most variable genes.

The 930 HI-vars are mostly rare alleles, with most being singletons or doubletons. There were only 93 variants with a frequency above 1% in the total joint called samples (Fig 6). Overall, the frequency distributions for sub-populations (SA, SC, FW and WE) are not uniform. The KS cluster (Khoe and San) is omitted due to low sample number. We note the dissimilarity when we consider the granularity of the data. In fact, some of the high impact variants tend to show a large disparity in frequency values between some clusters. For example, the *CYP27A1* rs114768494 variant (chr2:g.219677301C>T) (28th index in Fig 6) is only present in SC and WE with respective frequencies of 1.1% and 3.7%. Also, variants can exist in all the sub-populations but with significantly different proportions. For instance, the *CYP4B1* rs45446505 variant, (chr1:g.47279898C>T) (52nd index) is present at frequencies of 9.5%, 2.3%,4.5% and 3.5% for SA, SC, FW, and WE respectively. Another variant: *CYP4B1* rs3215983, (ch1:g.47280747_47280747del) (47th index) is common in the SC population with a 10% frequency. This value is at least twice that of other clusters. Frequency differences of ≈ 10% are observed in common high impact variants.

**Fig 6.**
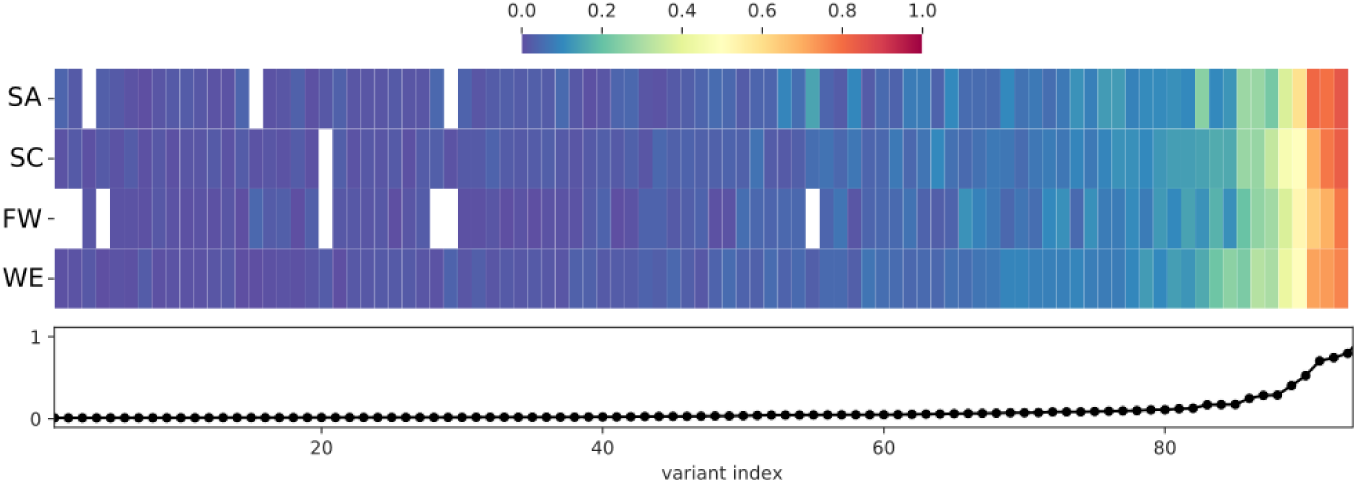
Distribution and frequency of HI-vars across sub-populations. Only common HI-vars (Frequency > 1% in the total joint called population dataset) are represented in the figure. The frequency per variant is reported in the lower panel of the figure as a line plot (indexed by frequency). The frequency of each of the variants in each sub-population is given by each heatmap column, with white indicating 0% frequency. See Table 2 for abbreviations

The regional overlap of the total HI-vars identified shows the majority of these variants are appear in one population cluster only (Fig 7). There are only ∼100 variants that overlap all African population clusters. These variants appearing in all regions have widely ranging frequencies, with most falling between 1-20% for the total African samples assessed. Each population cluster had >110 variants specific to it. Variants that occur only in one cluster are mostly rare, with an average frequency of less than 1% in their own respective cluster. Southern Africans have 20 cluster-specific variants with frequencies above 1% (20 variant) – more than any other cluster. Relatively fewer variants overlap between two clusters alone, with a trend of geographically close clusters sharing more variants than those which are distant.

**Fig 7.**
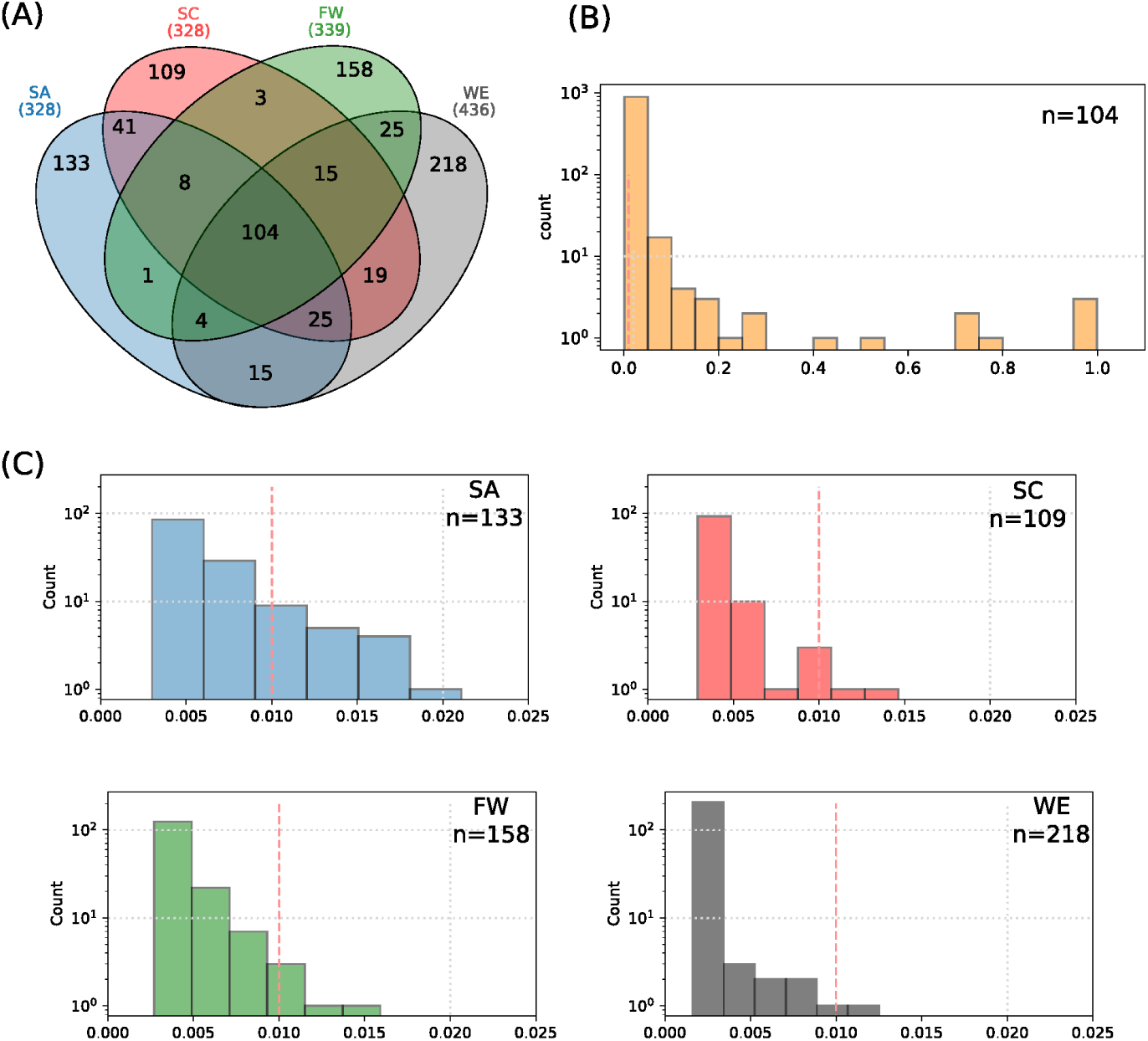
Characterisation of the HI-vars in clusters SA, SC, FW, and WE. (A) The Venn diagram shows the overlap between the clusters for these variants. Distribution plots of the frequencies of the common variants between the four clusters (B) and for the unique variants found in each sub-population (C).

Fixation index (*F*_*ST*_) assessments revealed that there are inter-cluster differences calculated for HI-vars (Figure 8 A), and also for all ADME gene variants (Fig 8 B). The greatest *F*_*ST*_ of all ADME variants is observed between SA and FW populations (0.0125) and the lowest *F*_*ST*_ is observed between SC and WE (0.003). For *F*_*ST*_ calculated across HI-vars, these are specific to ADME HI-vars as compared to HI-vars identified in random genes across the genome (n=2,000). This effect was significant between Far West Africans and all other clusters. Interestingly, despite being geographically close and having HI-vars in common, FW and WE clusters show an *F*_*ST*_ value of 0.0042, similar to the *F*_*ST*_ between the Far West FW and SC cluster, which are geographically distant and have no common variants – something meriting further study. Both of these differences show significant *p*-values of 9 × 10^−4^ and <10^−4^ between FW/WE and FW/SC respectively. *F*_*ST*_ values for all ADME gene variants overall show higher levels of differences, none of which, however, seem to be a property of these variants compared to genetic variants from a random set of genes (all *p*-values are non-significant).

**Fig 8.**
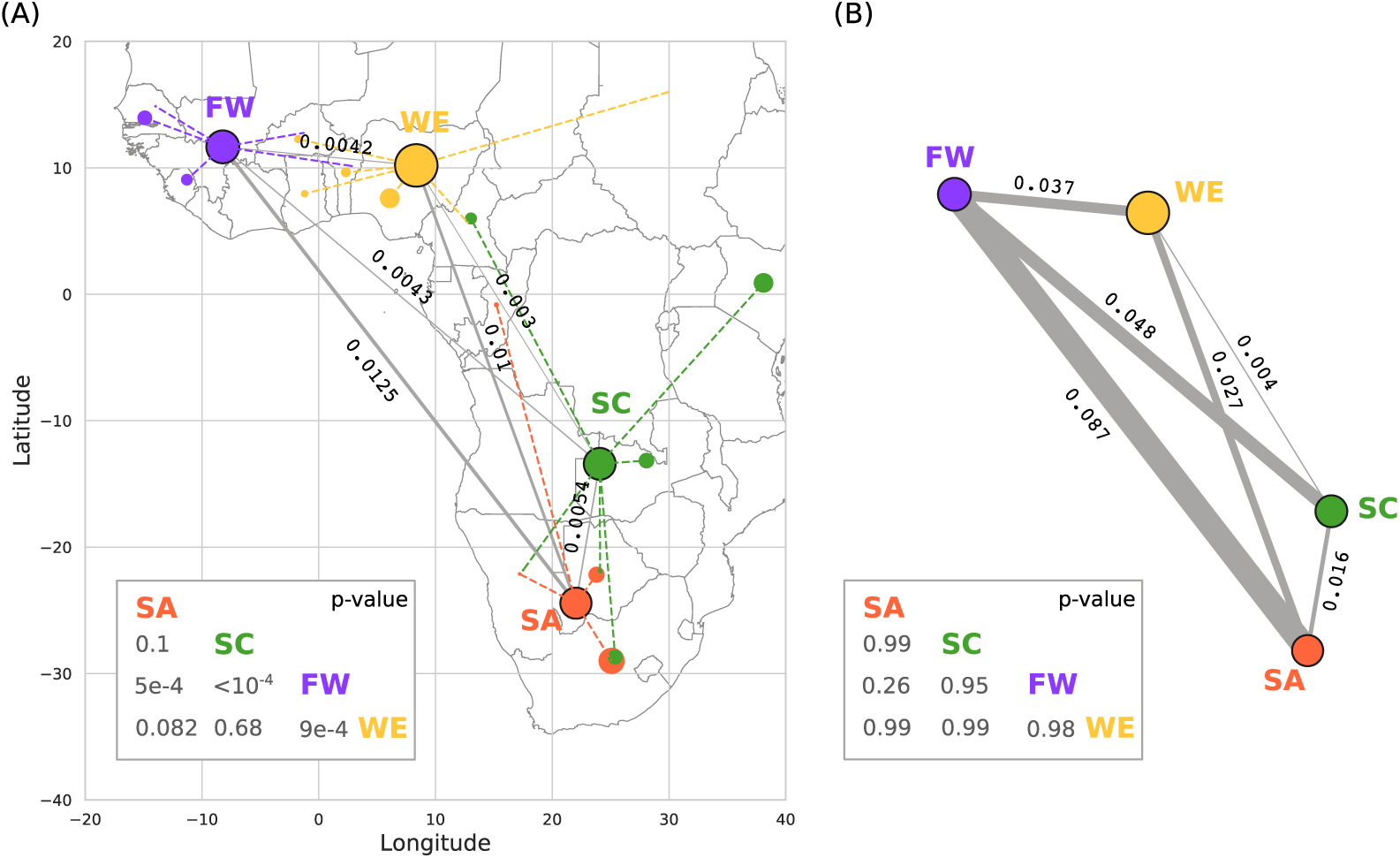
Calculation of Fixation index (*F*_*ST*_) between the population clusters. We used HI-vars (A) and all ADME variants (B) to compute the weighted *F*_*ST*_ value between each pair of populations using PLINK (version v1.90b6.3). The populations (SA-5) are represented by the geographical centroid of the ensemble of country centroids constituting each cluster. Dashed lines link the centroids of the countries to the cluster centroid. Node radii are proportional to the size of each sample. P-values were calculated from a random *F*_*ST*_ distribution by sampling 917 and 32,0983 variants of a random set of genes (n=2,000) for high impact variants and all ADME variants respectively.

In summary, we note that HI-vars are not uniform across African clusters, and that geographical proximity is not a proxy for genetic similarity in ADME genes.

### 2.5 CNVs

A copy number variant region (CNVR) is determined by aggregating overlapping CNVs identified in different individuals. A total of 259 CNVRs were identified, consisting of 106 duplications, 106 deletions and 47 mixed CNVRs (i.e. a region that is deleted in some individuals and duplicated in others) (Table 3). Duplications were further separated into biallelic duplications (3 or 4 copies) and multi-allelic duplications (> 4 copies). About 54% of CNVRs were unique, while the remaining CNVRs overlapped with one or more of the other CNVRs identified. Of the 299 ADME genes that were analysed a total of 116 genes (38.8%) contained at least one CNV. These include some important core pharmacogenes such as the *CYPs, UGTs* and *GTSs*. Furthermore, the number of CNVs in ADME genes per individual ranged from four to 71, with the majority of individuals (89.9%) harbouring between 11 and 30 CNVs.

**Table 3.** CNVs identified in core and extended ADME genes (percentages rounded to closest integer).

| CNV Category | Total | ADME genes |  |
| --- | --- | --- | --- |
|  |  | Core | Extended |
| Deletions | 106 (41%) | 30 | 76 |
| Biallelic duplications | 71 (27%) | 7 | 64 |
| Multi-allelic duplications | 35 (14%) | 2 | 33 |
| Mixed CNVs | 47 (18%) | 16 | 31 |
| Total | 259 | 55 (21%) | 204 (79%) |

### 2.6 Novel and highly differentiated variants

A novel variant in the context of this study is an SNV that is identified in the high coverage African population datasets, and not present in dbSNP (version 151) [13] which aggregates variants from various data sources that include the 1000 Genomes consortium [14, 15], GO-ESP [16], ExAC consortium [17], GnomAD [18] and TOPMED [19].

A total of 343,606 SNVs were called for the ADME genes from the HAAD set of 458 samples, with 12% classified as novel SNPs (Table 4). For the 32 core ADME genes, 5,818 novel variants were identified and a further 34,874 novel variants were identified in the 267 extended ADME genes within the HAAD. The majority of these variant types are intronic or intergenic variants (Fig 9). Of the novel coding variants, 8 were identified as HI-vars in core genes and 88 in extended genes.

**Table 4.** Summary of known and novel variants called from the HAAD for the ADME core and extended genes.

| HAAD ADME called datasets | Number of SNVs |
| --- | --- |
| Known ADME variants called | 304,666 |
| Novel ADME variants called | 40,692 |
| Core variants called – known | 44,039 |
| Core variants called – novel | 5,818 |
| Extended variants called – known | 260,627 |
| Extended variants called – novel | 34,874 |

**Fig 9.**
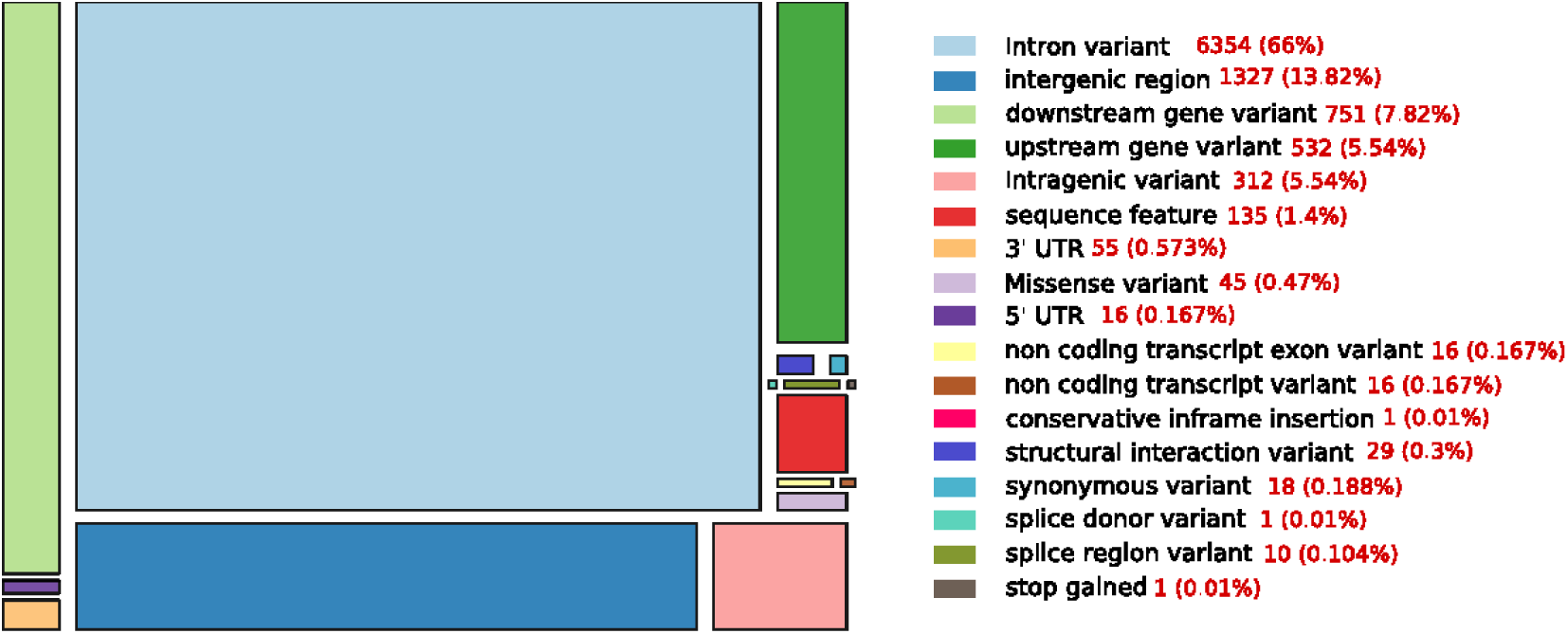
Distribution of ADME novel variant type by overall count.

The largest number of novel SNVs identified were from populations sampled from the Southern African region (not unexpected as there are no southern African populations in the KGP). Novel variants in each regional population cluster were characterised according to their effect as summarised in Table S1.

We compared the frequencies of ADME variants seen in the HAAD set as well as in at least one of the other large databases including 1000 Genomes Consortium, ExAC, gnomAD and TOPMED. Any variant with a frequency two-fold more or two-fold less in the HAAD set than in the other datasets was considered as highly differentiated. Approximately 1,957 ADME variants were highly differentiated in the HAAD data compared to 1000 Genomes consortium, ExAC, gnomAD and TOPMED datasets. Sixteen common variants with Minor Allele Frequency (MAF) ≥ 1% in eight core genes were more frequent in HAAD than in the KGP including African populations in those datasets One variant in one of the core genes (rs3017670, *SLC22A6*) was seen more commonly in the other datasets than in the HAAD data (Table S2). In total, 251 core and extended ADME genes harboured highly differentiated variants, with about 80% of them having at least 2 highly differentiated variants.

We performed a structural analysis of four rare novel HI-vars belonging respectively to *CYP2A13, CFTR, ABCB1*, and *NAT1* genes, all having an available protein structure from the Protein Data Bank (Fig 10). A variant chr19:g.41595975C>G causes a substitution p.Arg123Gly on *CYP2A13* (PDB code 2PG5) [20]. Mapping this variant on the structure shows a position close to the interaction site belonging to a rigid alpha helix which might affect the binding properties and the local folding integrity.

**Fig 10.**
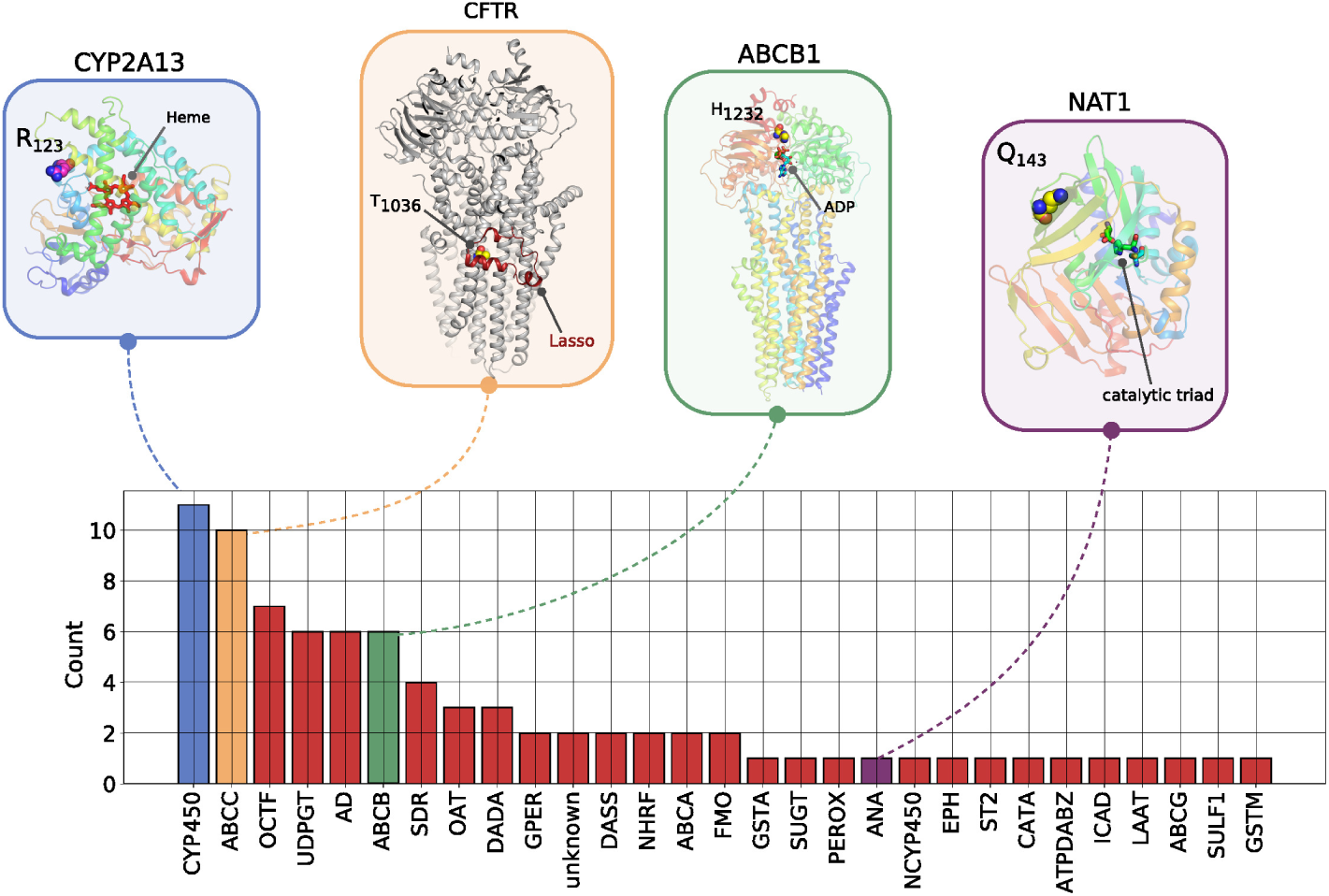
The number of novel high impact variants in ADME protein families. Abbreviation for Protein families: CYP450: cytochrome P450, OCTF: organic cation transporter family, UDPGT: UDP-glycosyltransferase, AD: aldehyde dehydrogenase, SDR: short-chain dehydrogenases/reductases, DADA: DAMOX/DASOX, GPER: glutathione peroxidase, DASS: SLC13A/DASS transporter, NHRF: nuclear hormone receptor family, GSTA: GST alpha, SUGT: Sugar transporter, PEROX: peroxidase, ANA: arylamine N-acetyltransferase, NCYP450: NADPH–cytochrome P450 reductase, EPH: Epoxide hydrolase, ST2: Sulfotransferase 2, CATA: cation transport ATPase, ATPDABZ: ATP-dependent AMP-binding enzyme, ICAD: iron-containing alcohol dehydrogenase, LAAT: L-type amino acid transporter, SULF1: sulfotransferase 1, GSTM: GST Mu. ABCC, ABCB, ABCA, FMO, ABCG are not abbreviated.

For the *CFTR* gene, a chr7:g.117250690A>G causes a substitution p.Thr1036A which is involved in the interaction of the Lasso domain of the protein serving as a critical interaction segment of CFTR with other proteins (PDB code 6MSM) [21, 22]. In addition, this threonine appears to form a pseudoproline-like structure in which the side chain OH is hydrogen bonded to its own backbone NH. This may contribute to the bending of the helix in which this residue is found. Mutation to Ala removes this hydrogen bond and may therefore influence the degree of bending of this helix.

A p.H1232Q protein variant in ABCB1 could affect the interactions of this residue with the ATP molecule required for the active transport process(PDB code 6C0V) [23]. In the structure His 1232 lies in a site to which an ATP is bound approximately 5 Åfrom the ATP gamma phosphate. Although not in direct contact with the ATP, it could interact with it via a network of hydrogen bonds involving water molecules or, if the histidine is protonated, via an electrostatic interaction with the ATP phosphates. A mutation to Gln could affect both types of interaction with the ATP.

The chr8:g.18079983C>T variant creates a premature stop codon in NAT1 gene (PDB code 2IJA). The variant corresponds to the position p.Q143 which is close to the catalytic site of the protein.

We analysed the distribution of the novel variants for the HAAD population cluster (Fig 11). The shared variants are generally exclusive for higher index values, which correspond to higher allele frequencies (Fig 11 A, B) in their respective cluster for both core (Fig 11 C) and extended genes (Fig 11 D). Moreover, we noted that the cluster specific variants cover a big portion of the frequency spectrum: most of them are rare (lower limit of the frequency spectrum).

**Fig 11.**
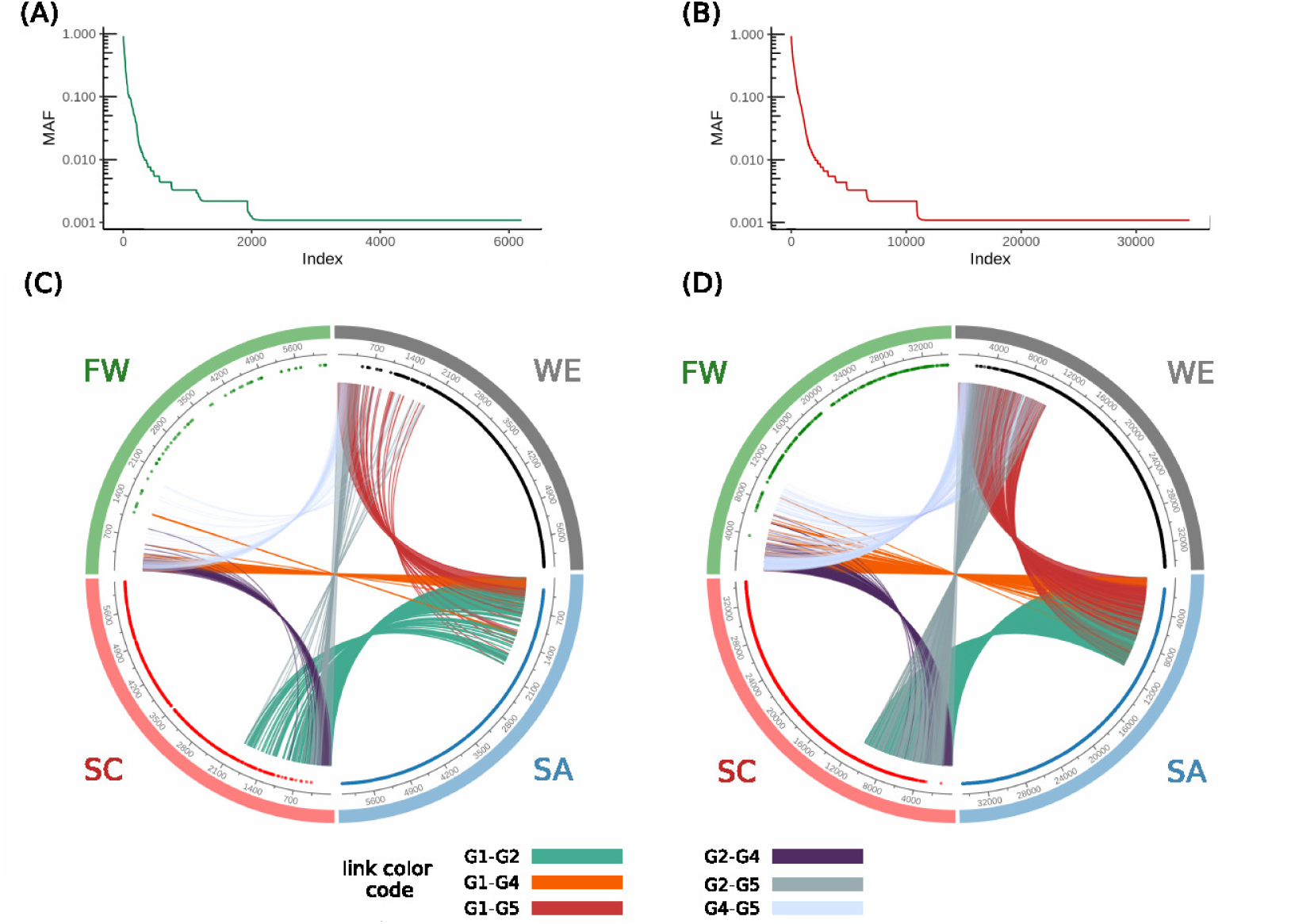
Characterisation of novel variant distribution across the ADME genes in HAAD. Variants are indexed by descending MAF for core genes (A) and extended genes (B) for the total population of HAAD samples. Circular plots for core (C) and extended genes (D) show the position of the unique variants per each cluster across the index axes represented by points. The index represents the MAF of the variant in the total HAAD dataset. A link is established if two clusters share the same variant. The FW cluster has fewer variants in common with other clusters in this figure due to low sample number used to generate this figure

### 2.7 Potential translational impact of ADME pharmacogenomic variants with known clinical effects

To assess the transferability of variants with known pharmacogenomic effect, we focused on variants with PharmGKB level 1A and 1B clinical annotations. A level 1A annotation denotes a variant-drug combination published as a CPIC guideline or known clinical implementation in a major health system, while a level 1B annotation denotes a variant-drug combination for which a large body of evidence shows an association in the context of altering drug response [24]. (Note that the absence of level 1 annotation may be evidence of lack of study of a variant, especially for African-specific variants, rather than evidence against clinical relevance.) In the entire HAAD set, we identified a total of 21 clinical variants (PharmGKB 1A/B) in 11 ADME genes. Nine of these variants had AF ≥ 0.05 in HAAD, while 12 are rarer (AF *<* 0.05). We next compared the frequency of the clinically actionable ADME gene variants in the combined HAAD population with that in the 1000 Genomes super populations as well as gnomAD (Table 5). Notably, two variants i.e. *CYP2D6*17* (rs28371706, AF = 0.2306) and the *CYP3A5*3* (rs776746, AF = 0.8315), had much higher frequencies in the African populations compared to the non-African KGP super populations as well as the combined gnomAD population. *CYP2D6*17* has been associated with decreased *CYP2D6* enzymatic activity in African Bantu populations [25].

**Table 5.**
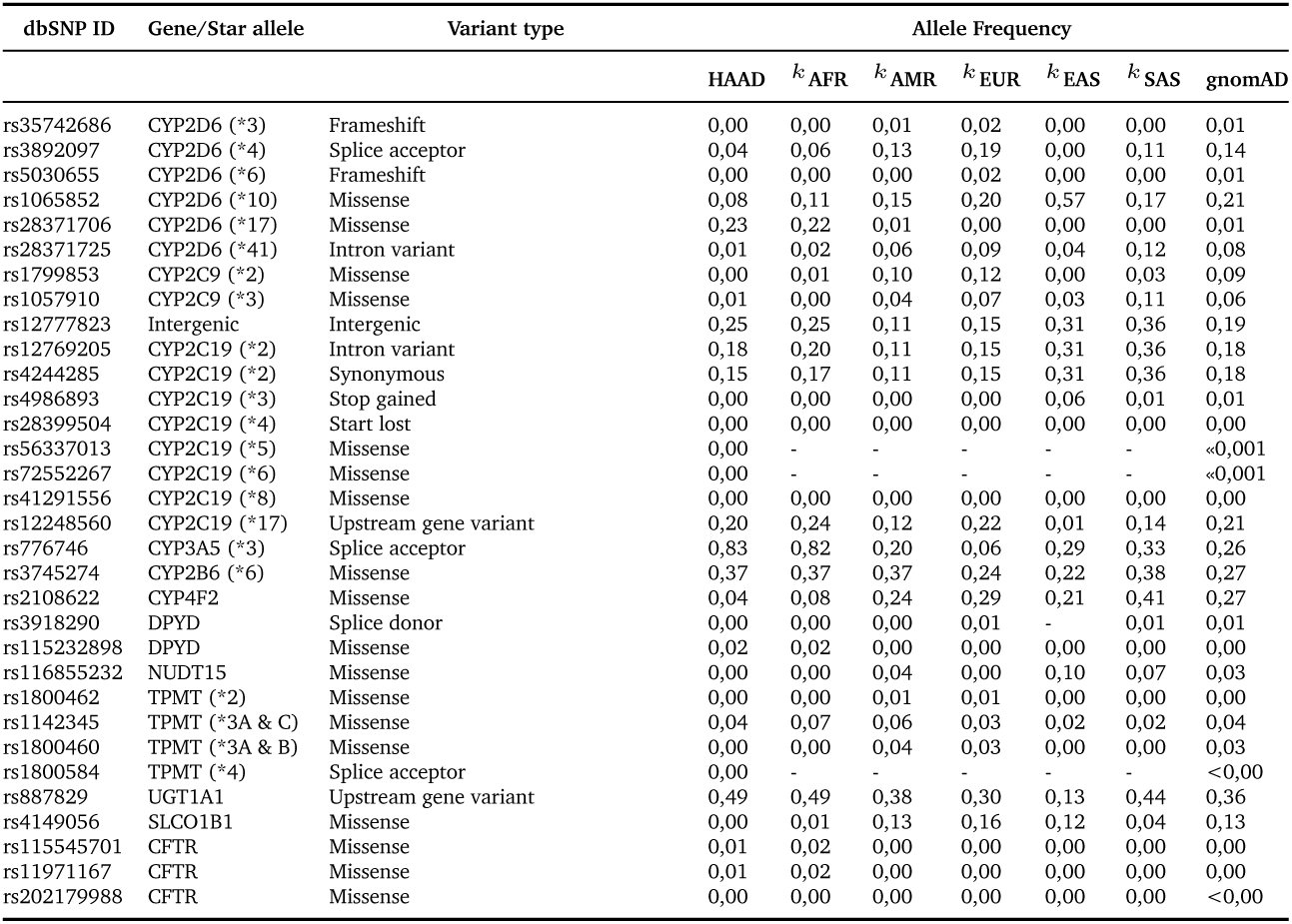
Allele frequency of the clinically actionable variants (PharmGKB 1A/B) in the combined HAAD dataset compared to the KGP super populations*k* as well as gnomAD. AFR: African, European: EUR, AMR: Ad Mixed American, EAS: East Asian, SAS: South Asian.

Some clinically-actionable ADME gene variants common in the non-African KGP super-populations are rare in the HAAD set. These include the variants *SLCO1B1* rs4149056 (*SLCO1B1*6*), *CYP4F2* rs2108622, *CYP2D6* rs3892097, *CYP2C9* rs1799853 and *CYP2C9* rs1057910. (Table 5).

Furthermore, we evaluated the distribution of level 1A/B PharmGKB variants within the African populations (HAAD and KGP) grouped according to the PCA clusters. Variants which show considerable frequency differences among clusters (SA, SC, FW, and WE) include *CYP2B6*6* (rs3745274), and *CYP2D6*17* (rs28371706) (Fig 12A). The number of level 1A/B PharmGKB variants per individual ranged from 0 to 15 (median of 6, 5, 5, and 6 in SA, SC, FW, and WE respectively) (Fig 12B), with 99.8% of individuals carrying at least one such variants.

**Fig 12.**
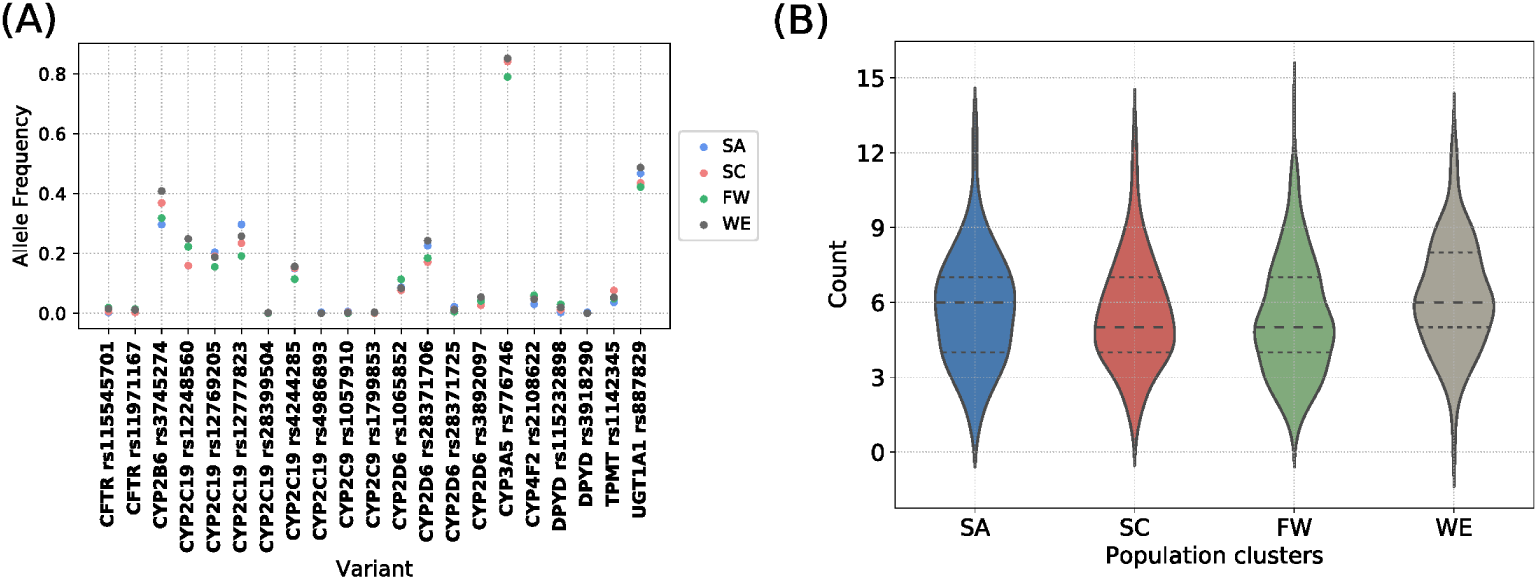
Distribution of pharmacogenomic variants with a high level of clinical annotation (PharmGKB level 1A/B). (A) Scatter plot of allele frequency of clinically relevant variants in the PCA clusters. (B) Violin plot of the number of clinically relevant variants carried per individual grouped by the population clusters.

### 2.8 Regulatory variation

There were 54 genetic variants across our African data sets in non-coding regions that have significantly higher prevalence than in the KGP overall data set (Table S3).

Fig S3 illustrates differences between population cluster pairs using *F*_*ST*_ scores. In most cases, the variability is not greater between pairs of population clusters (*F*_*ST*_ close to zero). We omit KS (Khoe and San) due to low sample size in this cluster.

### 2.9 Runs of homozygosity

Runs of homozygosity (ROH) are areas in the genome where an individual has two identical copies of the genome due to shared ancestors on the maternal and paternal lines. The size of the ROH correlates with how recent the shared ancestor was. With high coverage data, we are able to detect ROHs of at least 300kb in size. High ROH is a measure of inbreeding decreased fitness and may be associated with ill health [26, 27]. However, ROH are not randomly distributed across the genome and *islands of homozygosity* (ROHi) are known to exist: regions where the ROH of several individuals within a population overlap [28]. There is some evidence that these islands are found as a result of positive selection.

There are a total of 634 ROH in the sample. The key metrics we use are the size of ROHi (that is, how many individuals are in the ROHi) and size normalised by size of gene (ROHi/kb). The genes which have largest ROHi and ROHi/kb are *CYP1A1, CYP1A2*. The *ABCB1* and *DPYD* genes are relatively large genes and have a large ROHi. Tables S4 and S5 show a summary of the ROH found in the core and extended genes in our data sets. The range of ROHi/kb varies significantly across all genes in the genome Fig S4 shows a violin plot of the range of ROHi/kb in the core, extended, and all other genes in the genome. Statistical comparison is difficult because ranges are not normally distributed and a small number of extreme values skew the averages.

### 2.10 Coverage of ADME variants on SNV genotyping arrays

To evaluate whether genotyping array chips are suitable for detection of relevant ADME variants in African populations, we compared our whole genome sequencing variants with those captured by current arrays. Table 6 on page 21 shows the coverage of the variants that we detected in the core ADME genes in the WGS data compared to the Illumina Human Omni 2.5.8 (Omni) and the Illumina Infinium Multi-Ethnic AMR/AFR-8 Kit (MEGA). The Omni is a 2.39 million SNP array commonly used in human GWAS work – previous unpublished work shows that this is one of the best performing arrays on African populations. The MEGA array is 1.43 million SNP array optimised for African and Hispanic American populations (and can be augmented with approx. 200k user selected SNPs). For different minor allele frequencies of variants we detected (MAF) we show the number of variants that are at least at that threshold, the number of those variants captured by probes by the two arrays, and the percentage of the variants that are captured. As can be seen, even at relatively high frequencies, less than 5% of the variants are captured by the array for core genes, and less than 8% for extended genes. As expected the larger Omni does a better job. However, of the 93 common HI-vars, only 19 (20%) are on the Omni chip whereas 50 (54%) are on the MEGA.

**Table 6.**
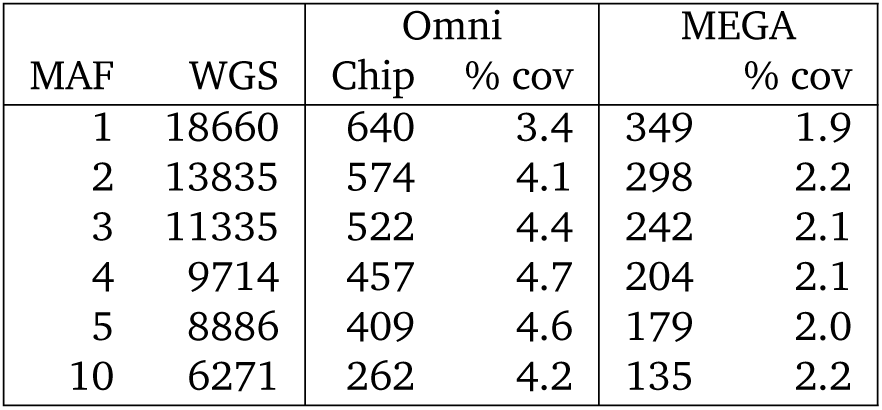
Variant coverage and overlap for core gene variants detected in HAAD whole genome sequencing datasets vs those captured by the Omni and the MEGA arrays. WGS=number of variants in the whole-genome data, Chip=number of variants in the chip, % cov= the percentage of SNPs at that MAF in WGS data that are covered by the array.

## 3 Discussion

Next generation technologies have provided pharmacogenomics and precision medicine a major increase in their application for disease treatment and drug safety [29]. ADME genes have been a focus due to their critical role in pharmacodynamics and pharmacokinetics. Our work presents the first study characterising the pharmacogenomics landscape of ADME genes in sub-Saharan Africa using high coverage whole-genome sequencing data which has been collected from different sources. The study’s main aim was to assess the variability of ADME genes across Africa and if this could have a significant impact on protein function and other pharmacologic properties and thus the potential impact on drug response.

We focus mainly on four African clusters distinguished geographically and genetically as shown by the PC whole-genome analysis. Overall assessments of structural and regulatory variation were evaluated across the complete dataset, while coding variants were assessed for functional impact. The applicability of known clinical variants and current genotyping technologies was also assessed.

In both novel variant and HI-vars analysis, our study demonstrates a significant level of variability. Most of the variants are rare and are population-specific in accordance with previous studies due mainly to increased population size and to a weak negative selection [24, 30–32]. Our high coverage data are adequate to genetically characterise these types of variants at high confidence levels. Evaluations of the false discovery rate of rare variants were previously estimated between 3.6% to 6.3% depending on the platform [33]. Therefore a broad extrapolation from our results is that there are between 30 to 60 false positive variants in our HI-vars. In the context of ADME pharmacogenes, although not all variants identified may prove to have functional impact, those that do may have significant consequences in dictating the drug-host response for individuals.

Our *F*_*ST*_ calculation highlights the differences between clusters. Calculation using all ADME variants led to values similar to results obtained for multiple sub-Saharan African ethnic groups that used 328,000 independent SNPs [34]. Genetic distance did not always correlate with geographical distance and in some pairs of clusters, the distance seems to be more significant in ADME genes. In the absence of clear evidence, it is not trivial to explain why two geographically close clusters like FW and WE, share a comparable degree of divergence like the pair FW-SC. Therefore, using ethno-geographical properties as a proxy to discriminate the pharmacogenomics landscape might be inaccurate.

In addition, the important number of cluster-specific novel and high impact rare variants suggest that strategies limited to studies of high-frequency alleles might be considered as an over-generalisation to a more complex pharmacogenomic landscape in sub-Saharan Africa. In fact, our work highlights a “genetic diversity bottleneck” for precision medicine applications, requiring a balance between variants useful for population-based applications (for a particular cluster of Africans) and between the potential impact posed by variants unique to the individual. Therefore, the complexities of variant interpretation and reporting in PGX testing [35] may be exacerbated by the complex African ADME landscape.

While some variants have similar frequencies in European and African populations, our assessment of the top-level clinically validated variants shows that these variants are more common in European populations than in African populations. This trend may be the result of the PGx knowledge bias towards European populations, with most variation in African and other global populations still largely uncharacterised in terms of PGx effect. Some variants show an opposite trend, such as the *CYP3A5*3* rs776746 and *CYP2D6*17* rs28371706, which are much more common in Africans than Europeans. These enzymes are known to be key metabolisers of a large number of drugs, and these two variants (as they are common) will impact the reliability of using a European based PGx strategy in African populations. Key drugs that may be affected by those variants are codeine [36], primaquine [7] (*CYP2D6*), and tacrolimus [37] (*CYP3A5*). We also see an interesting example of *SLCO1B1* rs4149056, which was seen in the KGP African populations (albeit rarely), which is not seen in the HAAD samples. This further reiterates the need for additional African sequences, as publicly accessible African genomic data cannot remain represented by the KGP alone. The greatest genomic coverage of African populations to date is available in genotyping array format [38]. These methods are unable to adequately characterise rare ADME variants at high confidence levels compared to high coverage WGS datasets. Moreover, we have also detected a large number of copy number variants, and were able to do so robustly with our high coverage sequencing data as compared to other methods [39]. The distribution of copy number variants and their impact on the ADME landscape in Africans is currently ongoing and will be available in a separate publication. As the state of data availability and type remains in flux, precision medicine approaches in Africa will be limited. In an ideal scenario, high coverage long read WGS will be used for more African samples undergoing clinical trials, as this allows for accurate resolution of haplotypes (including novel haplotypes), and thus, clearer interpretations of their potential impact on drug response.

## 4 Conclusion

Our work highlights that the ADME landscape in African populations is diverse, and shows the importance of rare variation held within individual population clusters. Therefore current array-based genotyping technologies have severe limitations to be applied as the high throughput method in precision medicine applications. As sequencing technology becomes more accessible and cheaper, characterisation of rare variants would benefit from the ongoing progress. Targeted sequencing and whole-exome sequencing would be better suited for characterising ADME genes. Moreover, a previous suggestion to consider intra-ethnic genetic characterisation in drug-development [8] might not be appropriate for sub-Saharan Africa due to the important presence of singletons and the subjective assigning of ethnicity for individuals The “genetic diversity bottleneck” in precision medicine might increase the burden of developing targeted therapies at sub-population levels because of the weak presence of common genetic patterns. However, these patterns might exist at the functional and phenotypic levels which might help to stratify the populations to clusters sharing common pharmacokinetic properties for a given drug. In this context, a proposed plan would integrate genotypic and phenotypic data into predictive models to unveil these patterns.

Capacity building efforts for pharmacogenetics and pharmacogenomics research in Africa is important. Strategies and policies for development of science and technology must ensure a future where Africa can take an active role in harnessing the power of genomic research in addressing its healthcare challenges. Promising positive steps are being taken with the establishment of initiatives such as the Human Heredity and Health in Africa project (http://h3africa.org/) that aims at strengthening research capacity for genomics in Africa.

### Limitations

There are many limitations of our work. The most obvious is the need for significantly more genomic data from Africa. Although, more samples are necessary generally, there is a particular need for more diverse sampling. We focus on sub-Saharan Africa, omitting northern Africa completely. We only had limited numbers of samples from Nilo-Saharan and Afroasiatic language speakers as well as speakers of non-Bantu languages in central, southern and eastern Africa (such as San and Khoe speakers). However, with more samples, we expect our conclusions to hold and the additional benefit would be a clearer resolution of the PGx landscape in diverse sub-clusters. Ideally, we would have merged the 1000 Genomes African data and the HAAD data set and done a combined analysis. However, the bulk of the 1000 Genomes WGS is low-coverage while the HAAD set is high-coverage which complicates comparative work significantly. As more data becomes available, this challenge will become easier. The discovery curve shown in Fig 5 shows we can expect to find many more variants when they are sequenced. Besides lack of genomic data, despite the effects of groups of excellence across Africa we have cited there is very little clinical and drug response data for African populations. Without this it will be difficult to associate the functional effect of variants to the clinical phenotypes. All of this costs money and requires scarce skills. Collaborations like ours, which has brought a diverse group of African scientists together show the potential of what can be done.

### Strengths

Our work investigates novel African datasets and combines these with established African sequences to assess as broad an overview of African ADME variation as possible. This work could lay the foundations for motivation of more PGx related studies in Africans. We applied diverse computational assessment methods to mine the data and retrieve valuable genomic information. This can assist in guiding future research in resource scarce environments.

## 5 Methods

### 5.1 Data

*H3A Consortium set* contains 272 samples selected and sequenced for the Human Heredity and Health in Africa (H3Africa) project. Samples cover populations from Benin, Burkina Faso, Botswana, Cameroon, Ghana, Nigeria and Zambia. Samples were shipped to the Human Genome Sequencing Center (HGSC) at Baylor College of Medicine (BCM), Houston, USA, under signed material transfer agreements from each project. Samples were prepared using the TruSeq Nano DNA Library Prep Kits and underwent whole genome sequencing on an Illumina TenX (150 bp) to a minimum depth of coverage of 30×.

*AWI-Gen set* consists of 100 South Eastern Bantu-Speakers (40× coverage).

*Cell Biology Research Unit, Wits set* consists of 40 samples from Soweto/Johannesburg South Africa (39 black and 1 mixed ancestry). Library preparation and sequencing was done at Edinburgh Genomics, Edinburgh, Scotland. Library preparation was done using the TruSeq Nano protocol and high coverage sequencing (∼30×) was done utilising the Illumina SeqLab workflow system and the Illumina HiSeqX platform.

The *SAHGP set* is a collection of 15 samples from the South African Human Genome Programme [11]. Two main Bantu-speaking ethno-linguistic groups were included: The Sotho (Sotho-Tswana speakers; n=8) and the Xhosa speakers (Nguni language; *n*=7 recruited from the Eastern Cape Province). The DNA samples were normalised to ∼60 ng/µl and ∼5 µg DNA was submitted to the Illumina Service Centre in San Diego, California, for sequencing on the Illumina HiSeq 2000 instrument (101 bp paired-end reads, ∼314 bp insert size) with a minimum read depth of coverage of 30× [11].

*SGDP set* contains 34 African samples selected from 300 individuals from the Simons Genome Diversity Project. Samples include populations from Congo, Namibia, Kenya, Senegal, Algeria, Nigeria, Gambia, Sudan and South Africa. Samples were sequenced at an average depth of 43× at Illumina Ltd; almost all samples were prepared using the same PCR-free library preparation [40].

*1000 Genome African set* consists of 507 African samples from 1000 Genomes Project. These samples include Gambian Mandinka, Mende from Sierra Leone, Yoruba from Ibadan, Nigeria, Esan from Nigeria and Luhya from Webuye, Kenya. Libraries were constructed on either Illumina HiSeq2000 or GAIIX with the use of 101 base pair end reads. Sequencing was done at an average depth of 4× [15].

The only phenotype made available to us was sex. In particular, self-identified ethnicity, location in the country, and disease status were not revealed.

### 5.2 Data processing

Table 8 shows the individual steps involved in creating the final joint called VCF of 966 samples.

**Table 7.**
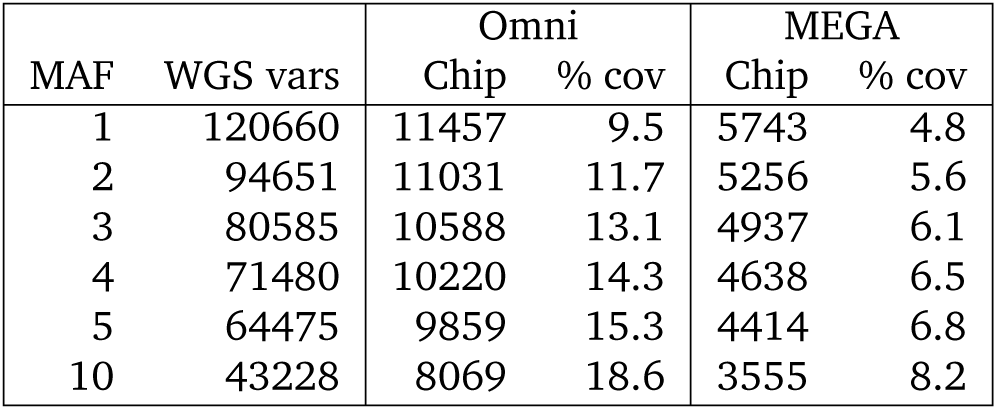
Variant coverage and overlap for extended gene variants detected in HAAD whole genome sequencing datasets vs those captured by the Illumnia Human Omni 2.5.8 array

**Table 8.**
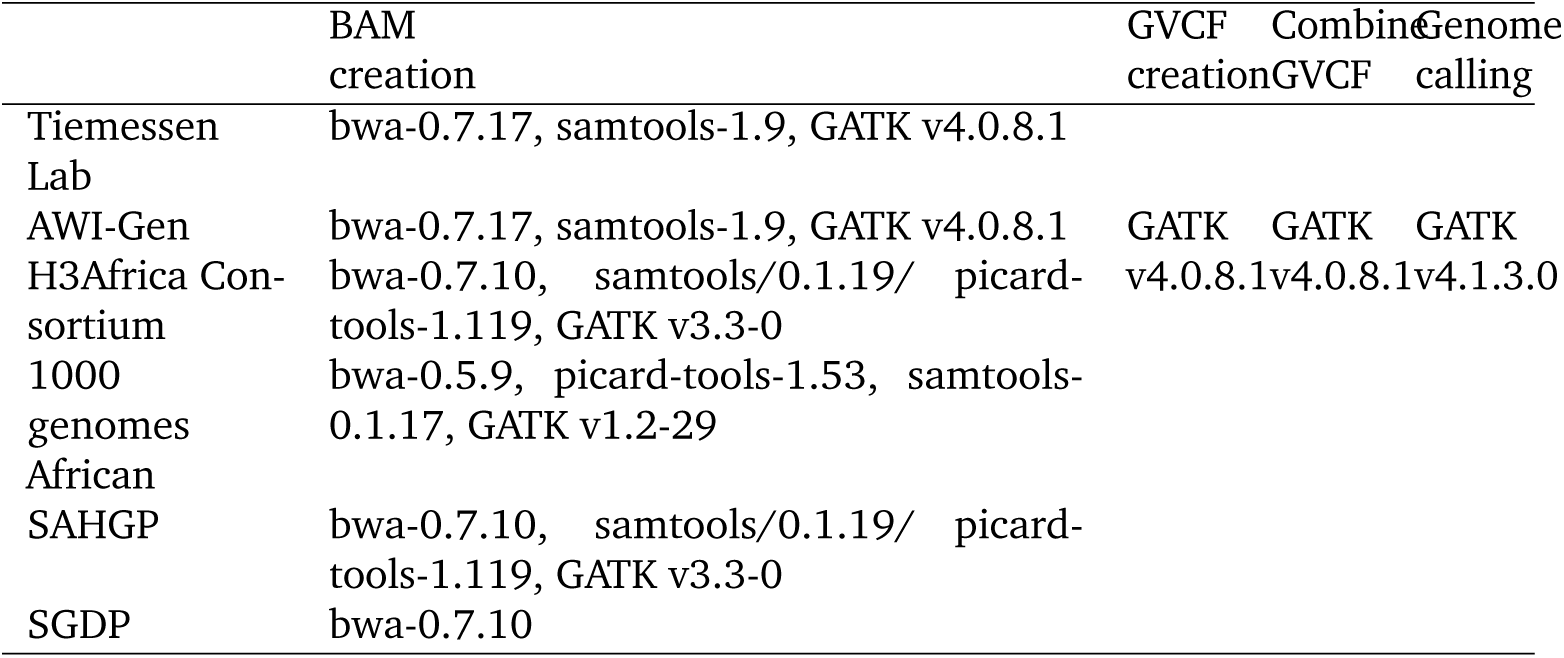
Processing done on individual samples and jointly. Tool versions are included

Most of the datasets mentioned had BAMs mapped against GRCh37 (also known as hs37d5) available. If BAMs were not available mapping was done from Fastqs. For all of the BAMs bwa-mem was used to do the alignment and Picard or GATK was used to MarkDuplicates and GATK was used to do Base Quality score recalibration for most of the cases. The only difference was the version of the specific tools being used in the alignment process.

From the BAMs we called gVCFs using HaplotypeCaller in gVCF mode using GATK v4.0.8.1. We combined all the gVCFs into one combined gVCF using GATK’s CombineGVCF (v4.0.8.1). From the combined gVCF we did joint calling using GenotypeGVCFs (v4.1.3.0) and followed GATK’s best practice for variant quality score recalibration for SNPs and INDELs. After applying VQSR we filtered for all the high quality (PASS) sites and used the VCF. The final VCF was used for downstream analysis. All code can be accessed at https://github.com/h3abionet/recalling.

### 5.3 Population structure

Population structure was computed using the autosomal data in our samples together with reference data sets in order to ensure a relatively unbiased structure. We included all 1000 Genomes Project African data, and two non-African 1000 Genomes Project sets (Utah residents (CEPH) with Northern and Western European ancestry – CEU – and Bengali in Bangladesh – BEB) and some chip data from various projects. Only unambiguous, biallelic SNPs (A/C, A/G, C/T, G/T) common in all data sets were used. The data was merged and pruned using PLINK [41], leaving 401k SNPs for analysis. Principal components were computed using PLINK and structure charts were produced using ADMIXTURE [42] (30 independent runs for each value of *k*) and averaged using CLUMPP [43]. All charts were produced with Genesis [44].

Population clusters were determined from the PCA values rather than from the project and self-identification labels due to overlapping data. The optimal number of clusters was determined using the method of Solovieff *et al*. [45], and clusters determined using *k*-means clustering with the R MASS package [46]. In analyses in which population clusters were compared, we only used the samples that appeared in the clusters (e.g., excluding Algerian, San samples). In all other analyses all the data was used. Choudury *et al*. [9] discusses the population structure of the H3A data in more detail.

### 5.4 ADME gene selection

ADME genes as defined by PharmADME (http://pharmaadme.org) (both core and extended definitions) were extracted using current genomic co-ordinates for GRCh37.p13, as obtained through BioMart [47]. Gene flanking regions were included in the extraction (10 000 bp upstream from gene start and downstream from gene end).

### 5.5 Annotation and Functional Prediction

Variants were classified and typed using SnpEff v4.3t [48] with the GR37Ch base reference for canonical gene transcripts. Variant Effect Predictor (VEP) v92.0 [49] was used for functional prediction based annotation. VEP was configured with dbNSFP v3.0 [50], a large database used to retrieve functional prediction scores for coding variants. The annotation analysis is implemented in g_miner workflow (https://github.com/hothman/PGx-Tools/tree/master/workflows/g_miner). An optimised model for functional prediction of pharmacogene variants produced by Zhou et al [51] was used as the basis for high impact classification of missense variants. The model uses five toolsets (LRT, MutationAssessor, PROVEAN, VEST3 and CADD). Loss of Function variants were classified as high impact if they were present in the canonical transcript of the gene. Singleton or doubleton high impact variants were filtered based on their VCF QUAL scores, using a cutoff of > 50. Any variant that did not match such criteria was removed prior to subsequent analyses with bcftools v1.9 [52]. Three HI-vars were not displayed in Fig 6 due to incorrect reference alleles inducing an erroneous frequency: ALDH3B1 rs11433668 and rs58160034; and ADH1C – rs283413. We have checked these variants in 1000 Genomes Project and gnomAD datasets to validate the error.

### 5.6 Fixation Index (*F*_*ST*_) analysis between population clusters

Differences between African subgroups were calculated by PLINK v1.9 [41], using mean, weighted *F*_*ST*_ between each pair of the population clusters. Prior to the calculation we applied linkage disequilibrium (LD) based pruning using PLINK v1.9 for different sets of variants: High Impact ADME, High Impact non-ADME, all ADME gene regions, and a set of 2000 random non ADME genes. The parameters used for this step are as follows: window size = 1000; step size = 5 and variance inflation factor = 2.

### 5.7 CNVs

Discovery and genotyping of CNVs was performed using GenomeSTRiP’s SVPreprocessing and CNVDiscovery (svtoolkit 2.00.1918) pipelines using the default parameters for genomes sequenced at 30-40× coverage [53].

### 5.8 Regulatory analysis

Genetic variants from ADME core genes were filtered for those meeting all the following criteria: in any non-coding region (10,000 bp up and downstream from canonical transcript); MAF >0.01; CADD-PHRED score ≥ 10 [54]; and binomial *p*-value compared with the entire 1000 Genomes Project data set < 0.05.

We compared these genetic variants (Table S3) for variability within pairs of populations as compared with the entire 1000 Genomes data set using *F*_*ST*_ scores [55] Since the number of genetic variants is small, we do not stratify it further into specific regulatory elements.

### 5.9 Runs of Homozygosity

Regions of homozygosity in core and extended gene sets were identified with using PLINK [41], using settings consistent for high-coverage data [26], viz. :–homozyg-snp 30:, :homozyg-kb 300:, :–homozyg-window-snp: :30:, :–homozyg group-verbose:.

## Ethics approval and consent to participate

No new data was generated specifically for this project – this is secondary analysis of data that had been generated and studied for other purposes. The H3A AWI-Gen Study (H3A data from Ghana, Burkina Faso and South Africa) was approved by the Human Research Ethics Committee (Medical) of the University of the Witwatersrand (Wits) (protocol numbers M121029 and M170880), and each contributing Centre obtained additional local ethics approval, as required. The H3A Benin study was approved by the Comité d’éthique de la recherche, Université de Montréal. The H3A CAfGEN study (Botswana) was approved by the IRB of the Ministry of Health of the Republic of Botswana (PPPME-13/18/1). The H3A TrypanoGEN Study (Cameroon component) was approved by the Comité National D’Ethique de la Recherche pour la Santé Humaine of the Republic of Cameroon (No 2013/11/364/L/CNERSH/SP). The H3A ACCME Study (Nigeria) was approved by the National Health Research Ethics Committee of Nigeria (NHREC/01/01/2007-29/11/2016). The H3A TrypanoGEN Study (Zambian component) was approved by the Biomedical Research Ethics Committee of the University of Zambia (FWA00000338). The data from the Cell Biology Research Lab, NICD/Wits was generated by a study approved by the Wits Human Research Ethics (Medical) Committee (protocol number M140926).

## Availability of data and materials

The data from the 1000 Genomes Project is publicly accessible from https://www.internationalgenome.org/data/. The Simons Genome Diversity Project data is available from EGA EGAS00001001959 and the Southern African Human Genome Project is available at EGAS00001002639 in EGA. The data from the Cell Biology Research Lab, NICD/Wits is available from Caroline Tiemessen on reasonable request, subject to ethics approval. The South African data from the AWI-Gen project is available from Michèle Ramsay on reasonable request (and will be deposited in EGA). All other datasets used in this study are available from the Human Heredity and Health in Africa (H3Africa) submission to EGA – EGAC00001000648.

## Authors’ contributions

JdR led the writing of the paper with assistance from HO. GB was responsible for the joint calling of the data and QC. JdR, HO, GB, SP, MM, SH contributed to the genomic data analysis. LC was primarily responsible for copy number variation analysis. DT and SA studied the transferability of ADME pharmacogenomic variants. PM provided the analysis of the regulatory regions. BD, DT, FMF, PM, MM, SP, GW, and SH all contributed to writing. SB, MC, CC, ASG, GS, CA, MoM, MR, GS, MS, CT and PT provided the critical analysis of the paper and contributed to writing. SH proposed the project, coordinated the work and co-led it with FJG. All authors contributed to writing the manuscript, and read and approved the final manuscript. This paper describes the views of the authors in their personal capacities and does not necessarily represent the official views of the funders.

## Acknowledgements

We thank the generous collaboration of the H3A AWI-Gen, CAfGEN, and TrypanoGEN group, the support of the Tiemessen Lab, the Awadalla Lab at the Université de Montréal, and Gabriel Anabawi from the University of Botswana. We thank Christine Clifton from GSK, Judith Kumuthini and Lyndon Zass for their advice and support, and Matthew Hall for his contributions to the project. This study would not have been possible without the generosity of the participants. We acknowledge the sterling contributions of our field workers, phlebotomists, laboratory scientists, administrators, data personnel and all other staff who contributed to the data and sample collections, processing, storage, and shipping. The AWI-Gen Collaborative Centre is funded by the NIH/NHGRI (Grant U54HG006938) as part of the H3Africa Consortium. MR is a South African Research Chair in Genomics and Bioinformatics of African Populations hosted by the University of the Witwatersrand, funded by the Department of Science and Technology, and administered by National Research Foundation of South Africa (NRF). The TrypanoGEN project was funded by the Wellcome Trust, study number 099310/Z/12/Z. The Collaborative African Genetics Network (CAfGEN) is funded by the NIH/NHGRI (grant CAfGEN 1U54AI110398). The whole genome sequencing of the H3A Data was supported by a grant from the National Human Genome Research Institute, National Institutes of Health (NIH/NHGRI) U54HG003273. Cell Biology Research Lab component: This work is based on the research supported by grants awards from the Strategic Health Innovation Partnerships (SHIP) Unit of the South African Medical Research Council, a grantee of the Bill & Melinda Gates Foundation, and the South African Research Chairs Initiative of the Department of Science and Technology and National Research Foundation of South Africa (84177). JdR was partially funded by the SA National Research Foundation (SFH170626244782). SP, GB and MM and the computational infrastructure used was partially supported by grants to the H3ABionet (NIH/NHGRI U41HG006941).

