## Supplementary Material for "The extent and impact of variation in ADME genes in sub-Saharan African populations"

**1** Sydney Brenner Institute for Molecular Bioscience, Faculty of Health Sciences, University of the Witwatersrand, Johannesburg, South Africa. **2** Division of Human Genetics, National Health Laboratory Service, and School of Pathology, Faculty of Health Sciences, University of the Witwatersrand, Johannesburg, South Africa. **3** Computational Biology Division and H3ABioNet, Department of Integrative Biomedical Sciences, University of Cape Town, South Africa. **4** Centre for Bioinformatics and Systems Biology, Faculty of Science, University of Khartoum, Sudan. **5** Department of Biochemistry & Medical Genetics, University of Manitoba **6** Department of Computer Science, Rhodes University, Makhanda, South Africa. **7** Neuroscience Research Program, Kleysen Institute for Advanced Medicine, Winnipeg Health Sciences Centre and Max Rady College of Medicine, University of Manitoba. **8** Department of Pharmacology and Therapeutics, Rady Faculty of Health Sciences, University of Manitoba, Winnipeg, Manitoba, Canada. **9** Institute for Human Virology, Abuja, Nigeria. **10** Institute of Human Virology and Greenebaum Comprehensive Cancer Center, University of Maryland School of Medicine, Baltimore, MD **11** Botswana-Baylor Children's Clinical Center of Excellence, Gaborone, Botswana. **12** Baylor College of Medicine, Houston, United States. **13** Molecular Parasitology and Entomology Unit, Department of Biochemistry, Faculty of Science, University of Dschang, Dschang, Cameroon. **14** Department of Disease Control, School of Veterinary Medicine, University of Zambia, Lusaka, Zambia. **15** Centre for HIV and STIs, National Institute for Communicable Diseases, National Health Laboratory Services and Faculty of Health Sciences, University of the Witwatersrand, Johannesburg South Africa. **16** Drug Metabolism & Pharmacokinetics, GlaxoSmithKline R&D, Ware, UK. **19** Data and Computational Sciences, GlaxoSmithKline R&D, Stevenage, UK. **17** Human Genetics, GlaxoSmithKline R&D, Stevenage, UK. **18** Clinical Pharmacology Modelling & Simulation, GlaxoSmithKline R&D, Sydney, Australia. **20** Global Health, GlaxoSmithKline R&D, Madrid, Spain. **21** School of Electrical & Information Engineering, University of the Witwatersrand, Johannesburg, South Africa. ‡Members of the Human Heredity and Health in Africa Consortium. Authors not marked with ‡ are members of the H3A/GSK ADME Collaboration but not of the H3Africa consortium.

### S1 Population Structure

A proper analysis of population structure is beyond the scope of this paper (See [1]). However it is useful to understand the extent of the diversity of the samples. Figure 2 in the main text showed a PCA of our samples (PC1 versus PC2). In Figure S1 we show PC2 versus PC3 of the same data, and we also show our data in the context of other populations. As explained in the methods section, the PC analysis included a number of reference populations including some

1000 Genomes European, Asian and African to ensure the analysis was unbiased. However, for clarity we only display some of the populations.

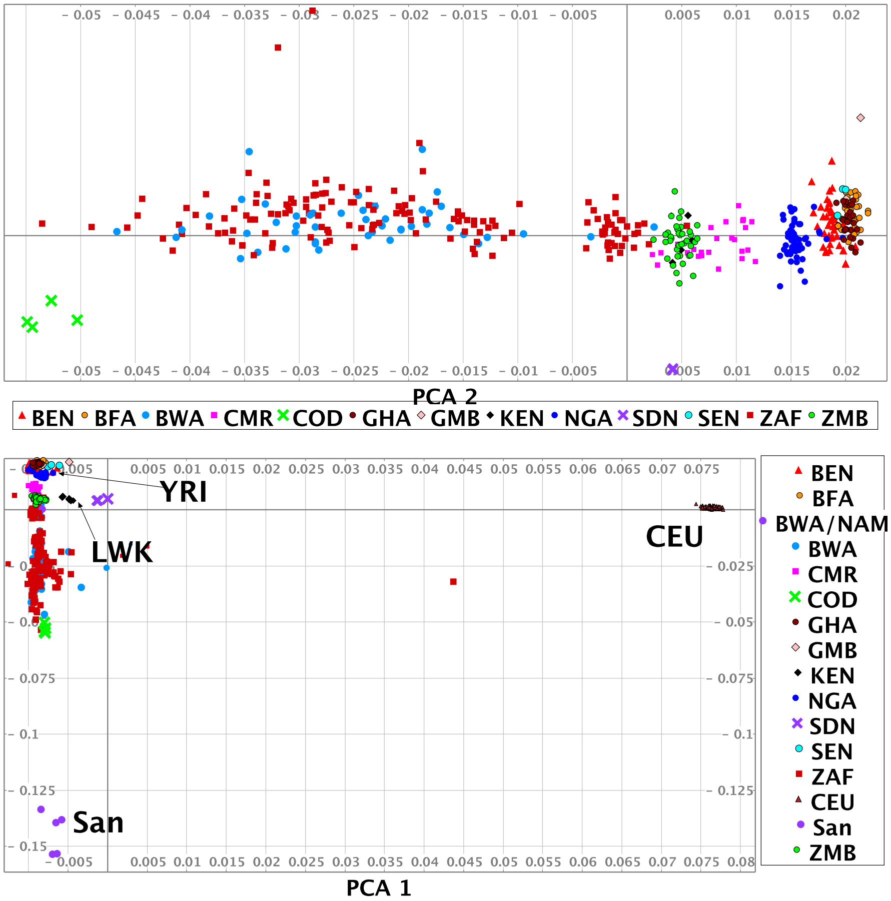


Figure S1: The figure on the top shows the PC2 versus PC2 of our data. The figure on the bottom shows PC1 versus PC2 – the same as Figure fig:sample-pca – in the context of other African and world populations.

Figure S2 shows the structure chart of the same data as that of Figure fig:sample-pca in the main paper, for $k = 3, \ldots, 8$. Admixture proportions were computed with ADMIXTURE [2] – 30 independent estimates were run for each value of $k$ and the final result computed using CLUMPP [3]. For clarity we have omitted some of the smaller groups. BWA=samples from Botswana, ZAF=samples from South Africa, NGA=Berom from Nigeria, BEN=Benin, CMR=Cameroon, BFA=Burkina Faso, GHA=Ghana, CEU=Utah residents (CEPH) with Northern and Western European ancestry (KGP), San=(Khoe and San from HAAD)

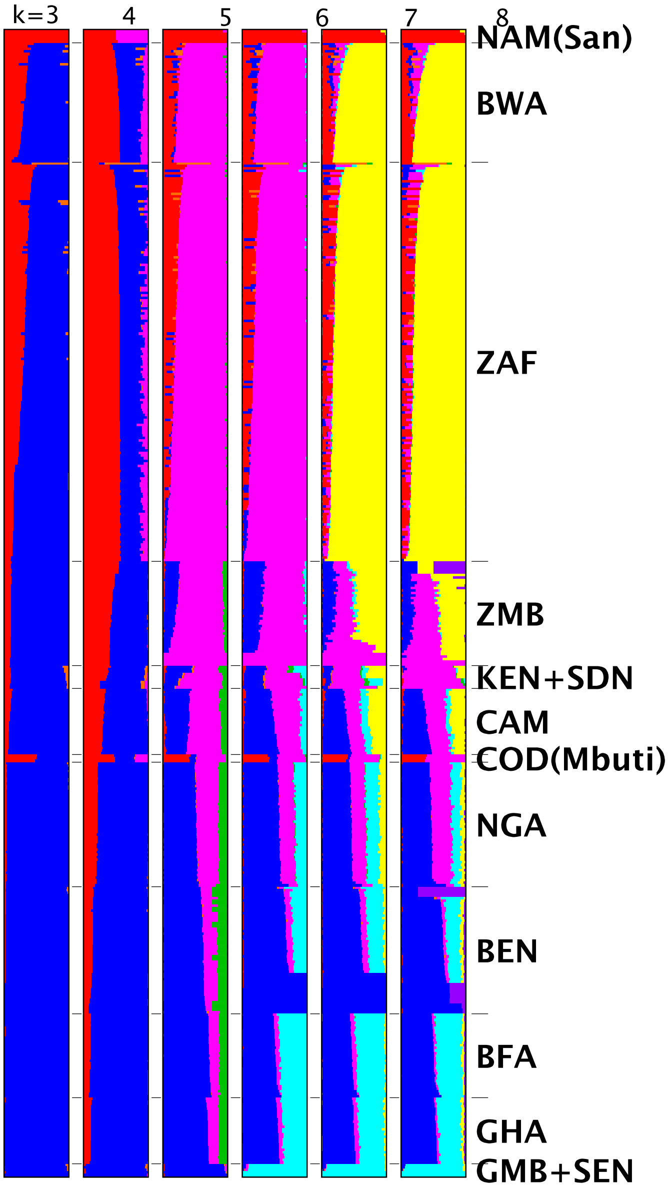


Figure S2: Structure chart – see text for description.

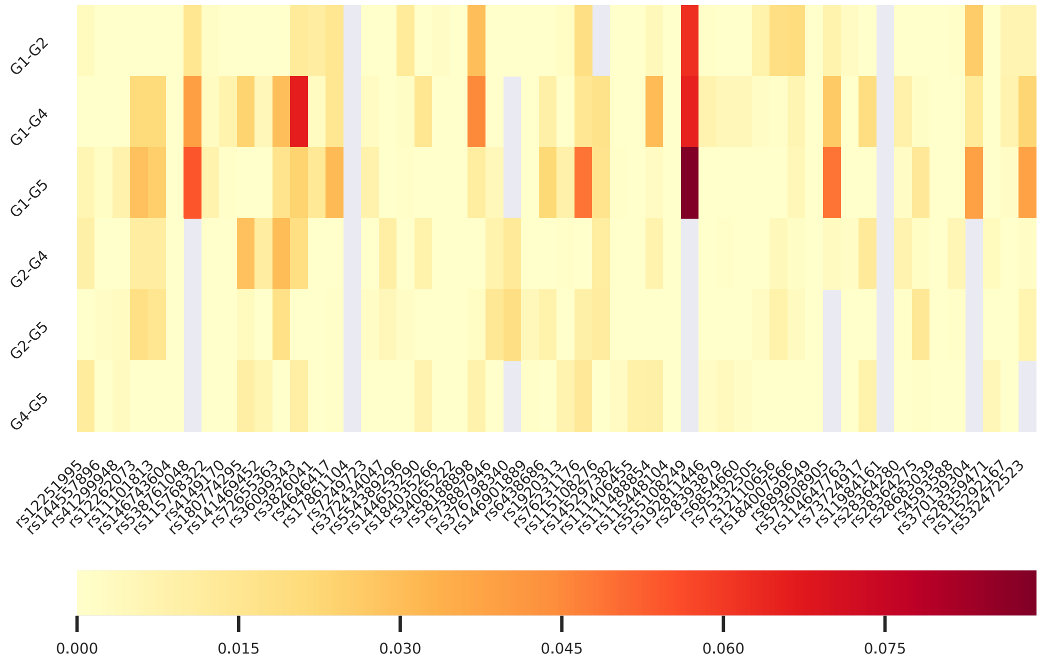


Figure S3: $F_{ST}$ scores for regulatory SNPs. The maximum value is $\approx 0.09$. Blanks are SNPs not scored between the given pairs.

#### S2 Regulatory Variation

Table S3 wlists SNPs after filtering for:

- in any non-coding region
- MAF $> 0.01$
- CADD-PHRED score $\geq 10$ [4]
- the canonical transcript for the SNP is not in a coding region (a check on the initial selection)
- binomial $p$-value compared with our entire 1000 Genomes data set $< 0.05$

The binomial $p$-value is calculated by taking the count of instances of the SNP as $2\times$homozygous count $+$ heterozygous count; the total is this count for both alleles and the expected probability is the MAF for the entire 1000 Genomes data set. Give that our data is generally high coverage and some of the 1000 Genomes data is not, this is a first cut at finding significant regulatory SNPs.

Some instances where the "ancestral" allele have much lower MAF than the "minor" allele pointing to a need to review calling the ancestral allele. We excluded examples with this issue (none in any case passed the $p$-value threshold).

This initial filtering resulted in 54 SNPs. We then compared differences between pairs of population versus the overall population by using PLINK to calculate $F_{ST}$ scores.

Table S1: Summary of known and novel variants called from the HAAD set for the ADME core and extended genes. Syn=Synonymous mutations; NS=non-synonymous mutations, BW/Nam = Sample from Botswana or Namibia

| | Categories of novel findings | | | | | | | | |
|---|---|---|---|---|---|---|---|---|---|
| | **variants** | **Rare** | **Indels** | **Single-tons** | **NS** | **Syn** | **Stop gained** | **Stop lost** | **LOF** |
| Algeria | 726 | 656 | 589 | 1002 | 1 | 0 | 0 | 0 | 0 |
| Benin | 4991 | 4074 | 2755 | 3260 | 31 | 13 | 1 | 0 | 0 |
| Botswana | 11274 | 8743 | 3492 | 8295 | 71 | 42 | 3 | 0 | 5 |
| BW/Nam | 1952 | 1364 | 1291 | 2090 | 4 | 2 | 0 | 0 | 0 |
| Burkina Faso | 5055 | 2343 | 2639 | 3807 | 26 | 19 | 1 | 0 | 1 |
| Cameroon | 4448 | 2358 | 2627 | 3421 | 31 | 15 | 2 | 0 | 2 |
| Congo | 2506 | 1363 | 1411 | 2336 | 12 | 5 | 0 | 0 | 1 |
| Gambia | 1110 | 1010 | 937 | 1390 | 0 | 1 | 0 | 0 | 0 |
| Ghana | 4527 | 2355 | 2585 | 3370 | 23 | 12 | 1 | 0 | 1 |
| Kenya | 2124 | 1496 | 1455 | 1896 | 9 | 2 | 0 | 0 | 0 |
| Namibia | 3591 | 1720 | 1864 | 3423 | 8 | 6 | 0 | 0 | 0 |
| Nigeria | 6916 | 6068 | 2927 | 5290 | 58 | 20 | 2 | 0 | 3 |
| Senegal | 1556 | 1268 | 1185 | 1543 | 5 | 1 | 0 | 0 | 0 |
| South Africa | 20492 | 18139 | 4523 | 13312 | 197 | 124 | 7 | 1 | 9 |
| Sudan | 1883 | 1181 | 1157 | 1883 | 4 | 6 | 0 | 0 | 0 |
| Zambia | 5296 | 2358 | 2768 | 3583 | 31 | 21 | 1 | 0 | 4 |

Table S2: Highly differentiated ADME variants in core genes in HAAD set. The frequency in different data sets is shown. Eff=Effect Effects: D=downstream; I=intron; M=missense; SR=splice region; SY=synonymous substitution; U=upstream

| **ID** | **Gene** | **Eff** | **AC** | **HAAD** | **KG** | **KG Afr** | **ExAC** | **ExAC/Afr** | **gnomAD** | **gno-mAD Afr** | **TOP MED** |
|---|---|---|---|---|---|---|---|---|---|---|---|
| rs113351578 | GSTP1 | SY | 31 | 0.0338 | 0.0 | 0.0 | 8.14e-06 | 0.0 | 3.23e-05 | 0.0 | 0.0 |
| rs112867476 | GSTT1 | M | 21 | 0.031 | 0.0 | 0.0 | 2.04e-05 | 8.94e-05 | 0.0 | 0.0 | 0.0 |
| rs1288697427 | SULT1A1 | SR | 11 | 0.012 | 0.0 | 0.0 | 7.92e-06 | 0.0 | 9.86e-05 | 0.000347 | 0.0 |
| rs187958013 | GSTT1 | SR | 23 | 0.0339 | 0.0002 | 0.0008 | 2.67e-05 | 0.000268 | 0.0 | 0.0 | 0.0 |
| rs1298375727 | ABCC2 | I | 10 | 0.0109 | 0.0 | 0.0 | 0.0 | 0.0 | 3.48e-05 | 0.0 | 0.0 |
| rs1303729583 | ABCB1 | U | 10 | 0.0109 | 0.0 | 0.0 | 0.0 | 0.0 | 3.24e-05 | 0.000115 | 0.0 |
| rs1468278213 | CYP2A6 | U | 10 | 0.0109 | 0.0 | 0.0 | 0.0 | 0.0 | 3.33e-05 | 0.000118 | 0.0 |
| rs111959737 | CYP2B6 | U | 13 | 0.0142 | 0.0 | 0.0 | 0.0 | 0.0 | 6.47e-05 | 0.00023 | 0.0 |
| rs111386238 | CYP2B6 | U | 13 | 0.0142 | 0.0 | 0.0 | 0.0 | 0.0 | 6.46e-05 | 0.00023 | 0.0 |
| rs112856484 | CYP2B6 | D | 12 | 0.0131 | 0.0 | 0.0 | 0.0 | 0.0 | 6.46e-05 | 0.000229 | 0.0 |
| rs368424346 | GSTT1 | SR | 13 | 0.0193 | 0.0 | 0.0 | 5.6e-05 | 0.000951 | 0.000213 | 0.000806 | 0.0 |
| rs376178963 | GSTT1 | U | 13 | 0.0193 | 0.0 | 0.0 | 5.87e-05 | 0.000965 | 0.0003 | 0.00113 | 0.0 |
| 10:101597866/CT | ABCC2 | I | 9 | 0.0107 | 0.0 | 0.0 | 0.0 | 0.0 | 0.000123 | 0.0 | 0.0 |
| rs990846266 | ABCC2 | I | 10 | 0.0109 | 0.0 | 0.0 | 0.0 | 0.0 | 0.000176 | 0.0 | 0.0 |
| rs1755884 | SLC22A2 | I | 593 | 0.353 | 0.0138 | 0.0015 | 0.0 | 0.0 | 0.0394 | 0.00866 | 0.0696 |
| rs3017670 | SLC22A6 | D | 891 | 0.0273 | 0.346 | 0.473 | 0.329 | 0.45 | 0.379 | 0.45 | 0.126 |

Table S3: Significant regulatory SNPs. Variations are labeled as the genomic coordinate, ancestral allele (A2), minor allele (A1). MAF for "All" is from 1000 Genomes; for African from our data as are homozygous and heterozygous counts; "missing" means there was no count for the allele in the at number of experiments. The *p*-value is the binomial *p*-value for our African MAF using the "All" fraction as the expected probability. Region: 3=3'UTR; 5=5'UTR; D=Downstream; NCE=non-coding exon; U=Upstream. A1=homozygous in minor allele; A2=homozygous in ancestral allele; het=heterozygous.

| | | | MAF | | Counts | | | | |
|---|---|---|---|---|---|---|---|---|---|
| Variation | Region | RS ID | All | African | A1 | het | A2 | missing | *p*-value |
| 8_18066300_G_C | U | rs28359471 | 0.0825 | 0.3319 | 43 | 218 | 197 | 0 | $1.1507 \times 10^{-101}$ |
| 11_62752289_C_T | 5 | rs4149170 | 0.2101 | 0.4083 | 76 | 222 | 160 | 0 | $5.3011 \times 10^{-42}$ |
| 4_89082070_G_A | U | rs555108249 | 0.0004 | 0.0284 | 0 | 26 | 432 | 0 | $5.6599 \times 10^{-39}$ |
| 3_121659860_C_T | 3 | rs1920313 | 0.0329 | 0.1321 | 8 | 105 | 345 | 0 | $1.0383 \times 10^{-37}$ |
| 15_75018218_G_A | U | rs36099343 | 0.0114 | 0.0764 | 4 | 62 | 392 | 0 | $8.2029 \times 10^{-35}$ |
| 3_121609955_A_G | D | rs6438686 | 0.0453 | 0.1496 | 14 | 109 | 335 | 0 | $5.7812 \times 10^{-34}$ |
| 4_69398821_T_A | D | rs145297382 | 0.0543 | 0.1731 | 35 | 65 | 290 | 68 | $2.2176 \times 10^{-32}$ |
| 7_87132251_T_G | D | rs11984161 | 0.1088 | 0.2424 | 25 | 172 | 261 | 0 | $2.5778 \times 10^{-30}$ |
| 10_96617356_C_T | D | rs538761048 | 0.0002 | 0.0197 | 0 | 18 | 440 | 0 | $6.0177 \times 10^{-30}$ |
| 22_42528538_G_A | D | rs58188898 | 0.0238 | 0.0987 | 11 | 67 | 373 | 7 | $9.1907 \times 10^{-29}$ |
| 12_21392205_C_T | D | rs72655363 | 0.0062 | 0.0524 | 2 | 44 | 412 | 0 | $1.8785 \times 10^{-28}$ |
| 6_160541549_T_A | U | rs12110656 | 0.0319 | 0.1114 | 7 | 88 | 363 | 0 | $4.4545 \times 10^{-27}$ |
| 6_160539330_T_A | U | rs75332505 | 0.0321 | 0.1103 | 7 | 87 | 364 | 0 | $2.7043 \times 10^{-26}$ |
| 7_99280641_G_A | U | rs28683039 | 0.0313 | 0.1081 | 3 | 93 | 362 | 0 | $6.0056 \times 10^{-26}$ |
| 15_75018890_A_C | U | rs3826041 | 0.4255 | 0.5941 | 165 | 213 | 79 | 1 | $9.8293 \times 10^{-25}$ |
| 15_75018931_G_C | U | rs4646417 | 0.0363 | 0.1146 | 5 | 95 | 358 | 0 | $1.5339 \times 10^{-24}$ |
| 10_101611798_T_G | 3 | rs12251995 | 0.0236 | 0.0895 | 3 | 76 | 379 | 0 | $4.2030 \times 10^{-24}$ |
| 4_89083634_A_G | U | rs192811446 | 0.004 | 0.0393 | 0 | 36 | 422 | 0 | $8.7280 \times 10^{-24}$ |
| 4_89084896_G_T | NCE | rs6854660 | 0.004 | 0.0393 | 0 | 36 | 422 | 0 | $8.7280 \times 10^{-24}$ |
| 10_135355400_C_T | D | rs12262073 | 0.0421 | 0.1212 | 2 | 107 | 349 | 0 | $8.4664 \times 10^{-23}$ |
| 10_135355411_C_A | D | rs11101813 | 0.0421 | 0.1201 | 2 | 106 | 350 | 0 | $2.6654 \times 10^{-22}$ |
| 7_87133538_A_G | 3 | rs28364275 | 0.0218 | 0.0797 | 6 | 61 | 391 | 0 | $1.1276 \times 10^{-20}$ |
| 6_160542819_T_C | U | rs6899549 | 0.0192 | 0.0742 | 4 | 60 | 394 | 0 | $1.4139 \times 10^{-20}$ |
| 6_18126648_C_G | D | rs114647763 | 0.0268 | 0.0841 | 4 | 69 | 385 | 0 | $4.9648 \times 10^{-18}$ |
| 8_18247940_A_G | U | rs532472523 | 0.001 | 0.0186 | 0 | 17 | 441 | 0 | $2.3327 \times 10^{-16}$ |
| 3_121665481_C_T | D | rs76231176 | 0.0188 | 0.0611 | 3 | 50 | 405 | 0 | $4.7880 \times 10^{-14}$ |
| 4_69957763_C_T | U | rs111488854 | 0.007 | 0.0360 | 2 | 29 | 427 | 0 | $6.7940 \times 10^{-14}$ |
| 6_18124135_G_A | D | rs573608905 | 0.0012 | 0.0175 | 0 | 16 | 442 | 0 | $6.8944 \times 10^{-14}$ |
| 10_101612267_T_C | D | rs144557896 | 0.0066 | 0.0349 | 1 | 30 | 427 | 0 | $7.7669 \times 10^{-14}$ |
| 15_75018932_G_A | U | rs17861104 | 0.01 | 0.0426 | 0 | 39 | 419 | 0 | $1.3425 \times 10^{-13}$ |
| 11_67348925_A_T | U | rs180774295 | 0.0032 | 0.0229 | 0 | 21 | 437 | 0 | $6.5138 \times 10^{-12}$ |
| 22_42531473_A_G | D | rs376798340 | 0.0004 | 0.0110 | 1 | 8 | 446 | 3 | $7.7221 \times 10^{-12}$ |
| 8_18065932_G_A | U | rs370139304 | 0.0006 | 0.0120 | 0 | 11 | 447 | 0 | $1.9831 \times 10^{-11}$ |
| 19_41348142_T_A | D | rs372434047 | 0.0026 | 0.0197 | 0 | 18 | 440 | 0 | $8.8238 \times 10^{-11}$ |
| 8_18063651_T_G | U | rs4593588 | 0.0116 | 0.0371 | 1 | 32 | 425 | 0 | $6.8774 \times 10^{-09}$ |
| 3_121609253_G_A | D | rs146901889 | 0.0178 | 0.0459 | 1 | 40 | 417 | 0 | $5.4202 \times 10^{-08}$ |

| | | | | | | | | | |
|---|---|---|---|---|---|---|---|---|---|
| 10_96834043_C_T | U | rs115768322 | 0.004 | 0.0197 | 0 | 18 | 440 | 0 | $6.2949 \times 10^{-08}$ |
| 4_89007744_T_A | D | rs115448104 | 0.0104 | 0.0317 | 0 | 29 | 429 | 0 | $2.3959 \times 10^{-07}$ |
| 10_135354531_C_A | D | rs41299948 | 0.0136 | 0.0371 | 0 | 34 | 424 | 0 | $2.7682 \times 10^{-07}$ |
| 10_96616487_T_A | D | rs146743604 | 0.0084 | 0.0273 | 1 | 23 | 434 | 0 | $5.0394 \times 10^{-07}$ |
| 6_160541919_T_C | U | rs184007566 | 0.0042 | 0.0175 | 1 | 14 | 443 | 0 | $2.8005 \times 10^{-06}$ |
| 6_18156129_G_A | 5 | rs73724917 | 0.0028 | 0.0142 | 0 | 13 | 445 | 0 | $2.9653 \times 10^{-06}$ |
| 1_98386622_G_A | U | rs554389296 | 0.006 | 0.0208 | 0 | 19 | 438 | 1 | $4.6863 \times 10^{-06}$ |
| 22_42529156_C_T | D | rs73887946 | 0.0074 | 0.0200 | 1 | 16 | 434 | 7 | $1.9377 \times 10^{-04}$ |
| 22_24373140_G_T | D | rs184035266 | 0.0036 | 0.0147 | 3 | 4 | 333 | 118 | $2.2664 \times 10^{-04}$ |
| 19_41345991_G_A | D | rs7249723 | 0.0573 | 0.0841 | 7 | 63 | 388 | 0 | $6.2582 \times 10^{-04}$ |
| 7_87133243_C_T | 3 | rs28364280 | 0.0084 | 0.0197 | 1 | 16 | 441 | 0 | $9.8999 \times 10^{-04}$ |
| 4_89084520_G_T | U | rs28393879 | 0.0064 | 0.0164 | 0 | 15 | 443 | 0 | $1.0730 \times 10^{-03}$ |
| 2_234686908_A_G | D | rs34065222 | 0.01 | 0.0218 | 0 | 20 | 438 | 0 | $1.2168 \times 10^{-03}$ |
| 8_18084765_T_G | D | rs115292167 | 0.006 | 0.0131 | 0 | 12 | 446 | 0 | $1.0697 \times 10^{-02}$ |
| 4_69402508_A_G | D | rs111406455 | 0.0128 | 0.0230 | 3 | 12 | 376 | 67 | $1.3828 \times 10^{-02}$ |
| 1_98389412_T_A | U | rs144653290 | 0.0056 | 0.0120 | 0 | 11 | 447 | 0 | $1.5921 \times 10^{-02}$ |
| 3_121665494_T_G | D | rs115108276 | 0.0054 | 0.0109 | 0 | 10 | 448 | 0 | $2.9545 \times 10^{-02}$ |
| 12_21070798_G_A | D | rs141469452 | 0.007 | 0.0131 | 0 | 12 | 446 | 0 | $3.0569 \times 10^{-02}$ |

Some notes on most statistically-significant variants:

- rs28359471 (`http://grch37.ensembl.org/Homo_sapiens/Variation/Explore?db=core;r=8:18065800-18066800;v=rs28359471;vdb=variation;vf=477979426`) – NAT1 gene, includes two intronic transcripts and binding sites for two TFs: E2F1::ELK1 and ETV2::RFX5; gene expression correlations for:
  - Thyroid, Skin Sun Exposed Lower leg, Muscle Skeletal, Heart Atrial Appendage, Thyroid, Nerve Tibial, Artery Tibial
- rs4149170 (`http://grch37.ensembl.org/Homo_sapiens/Variation/Population?db=core;r=11:62751789-62752789;v=rs4149170;vdb=variation;vf=147699`) – SLC22A6 gene, includes three 5′ UTR and one NMD variant and binding sites for TFs: GCM1::CEBPB, GCM1::NHLH1, GCM1::SOX2, GCM2::PITX1, MAX, TFAP4::MAX, CLOCK::BHLHA15, SPDEF, GCM1::ETV7, GCM1::SPDEF, HOXB2::RFX5, TEAD4::CEBPD, TEAD4::RFX5, TFAP2C::MAX, E2F1::ELK1, ETV2::NHLH1, ETV2::ONECUT2, ERF::ONECUT2, FLI1::ONECUT2; gene expression correlations for:
  - Brain Hippocampus, Pituitary, Brain Cortex, Artery Aorta, Uterus, Artery Coronary, Brain Anterior cingulate cortex BA24, Brain Putamen basal ganglia
- rs555108249 (`http://grch37.ensembl.org/Homo_sapiens/Variation/Explore?db=core;r=4:89081570-89082570;v=rs555108249;vdb=variation;vf=517892906`) – ABCG2 gene, includes 2 intronic transcripts, no reported regulatory features
- rs1920313 (`http://grch37.ensembl.org/Homo_sapiens/Variation/Explore?db=core;r=3:121659360-121660360;v=rs1920313;vdb=variation;vf=323181359`) – ABCG2 gene, includes 2 3′ UTR variants; gene expression correlations for:
  - Whole Blood, Cells Transformed fibroblasts

- rs36099343 (`http://grch37.ensembl.org/Homo_sapiens/Variation/Explore?db=core;r=15:75017718-75018718;v=rs36099343;vdb=variation;vf=445364005`) – CYP1A1 gene, TF binding site ETV2::GSC2
  - Whole Blood

These examples indicate considerable regulatory variation in ADME SNPs and hence point to more complex analysis in future work.

#### S3 Runs of Homozygosity

Figure S4 shows the distribution of runs of homozygosity across the genes, showing the density of ROHi per gene, normalised by gene length

Figure S4: Distribution of the ROHi/kb across the core, extended, and all genes. As there are extreme values, a $y$-cut-off of 3 was chosen to assist comparison. The median value and inter-quartile range is shown.

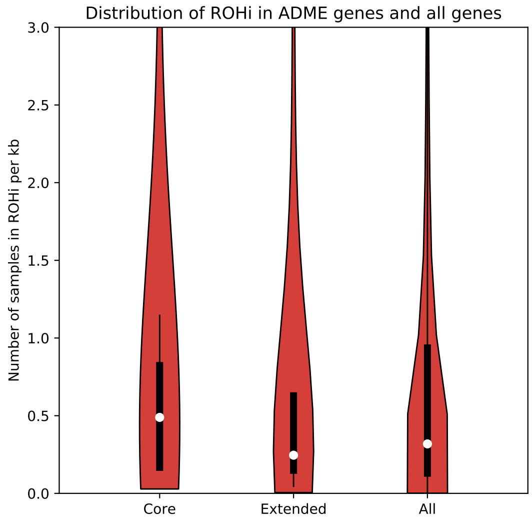


Table S4 shows the ROHi found in the core genes. For each group, the proportion of the individuals that are part of that ROHi for that gene is shown. In the two rightmost columns, the total number of individuals in the data set that are part of the ROHi is shown and then that number normalised by the length of the gene (i.e., #ROHi per thousand base pairs). In Table S5 a similar table is given for the extended data set.

Table S4: Runs of homozygosity in the core genes split by group. For each gene the number of ROH found across all samples is shown by group as a fraction of the individuals in that group who share the ROHi, followed the total number and the total normalised by gene length.

| **Gene** | **SA** | **SC** | **KS** | **FW** | **WE** | **O** | # **ROHi** | **ROHi/kb** |
|---|---|---|---|---|---|---|---|---|
| ABCB1 | 0.33 | 0.27 | 0.33 | 0.25 | 0.41 | 0.00 | 98 | 0.47 |
| ABCC2 | 0.04 | 0.03 | 0.00 | 0.00 | 0.04 | 0.00 | 10 | 0.14 |
| ABCG2 | 0.02 | 0.02 | 0.00 | 0.00 | 0.01 | 0.00 | 4 | 0.03 |
| CYP1A1 | 0.17 | 0.30 | 0.00 | 0.00 | 0.24 | 0.50 | 64 | 10.68 |
| CYP1A2 | 0.12 | 0.30 | 0.00 | 0.00 | 0.12 | 0.50 | 46 | 5.93 |
| CYP2A6 | 0.00 | 0.00 | 0.00 | 0.00 | 0.00 | 0.50 | 1 | 0.14 |
| CYP2B6 | 0.00 | 0.00 | 0.00 | 0.00 | 0.00 | 0.50 | 1 | 0.04 |
| CYP2C19 | 0.17 | 0.10 | 0.00 | 0.25 | 0.16 | 0.00 | 43 | 0.48 |
| CYP2C8 | 0.08 | 0.07 | 0.00 | 0.00 | 0.08 | 0.00 | 21 | 0.64 |
| CYP2C9 | 0.10 | 0.10 | 0.00 | 0.25 | 0.12 | 0.00 | 31 | 0.61 |
| CYP2D6 | 0.04 | 0.03 | 0.00 | 0.00 | 0.04 | 0.00 | 11 | 2.51 |
| CYP2E1 | 0.01 | 0.00 | 0.00 | 0.00 | 0.00 | 0.00 | 1 | 0.09 |
| CYP3A4 | 0.09 | 0.10 | 0.33 | 0.00 | 0.06 | 0.00 | 23 | 0.84 |
| CYP3A5 | 0.09 | 0.10 | 0.33 | 0.25 | 0.09 | 0.00 | 27 | 0.85 |
| DPYD | 0.15 | 0.28 | 0.00 | 0.50 | 0.38 | 0.00 | 78 | 0.09 |
| DPYD-AS1 | 0.01 | 0.03 | 0.00 | 0.00 | 0.03 | 0.00 | 6 | 0.03 |
| DPYD-AS2 | 0.13 | 0.27 | 0.00 | 0.50 | 0.32 | 0.00 | 68 | 60.12 |
| GSTM1 | 0.00 | 0.05 | 0.00 | 0.00 | 0.01 | 0.00 | 4 | 0.67 |
| GSTP1 | 0.06 | 0.02 | 0.00 | 0.25 | 0.05 | 0.00 | 14 | 4.58 |
| GSTT1 | 0.03 | 0.03 | 0.00 | 0.00 | 0.02 | 0.00 | 7 | 0.86 |
| NAT1 | 0.03 | 0.08 | 0.33 | 0.00 | 0.00 | 0.00 | 9 | 0.17 |
| NAT2 | 0.03 | 0.05 | 0.00 | 0.00 | 0.00 | 0.00 | 6 | 0.60 |
| SLC15A2 | 0.12 | 0.05 | 0.00 | 0.00 | 0.09 | 0.00 | 25 | 0.50 |
| SLC22A1 | 0.01 | 0.02 | 0.00 | 0.00 | 0.02 | 0.00 | 4 | 0.11 |
| SLC22A2 | 0.00 | 0.02 | 0.00 | 0.00 | 0.02 | 0.00 | 3 | 0.07 |
| SLC22A6 | 0.07 | 0.00 | 0.00 | 0.00 | 0.03 | 0.00 | 10 | 1.19 |
| SLCO1B1 | 0.07 | 0.08 | 0.00 | 0.00 | 0.05 | 0.00 | 18 | 0.17 |
| SLCO1B3 | 0.06 | 0.10 | 0.00 | 0.00 | 0.05 | 0.00 | 18 | 0.17 |
| SULT1A1 | 0.00 | 0.00 | 0.00 | 0.00 | 0.00 | 0.50 | 1 | 0.06 |
| TPMT | 0.02 | 0.03 | 0.00 | 0.00 | 0.02 | 0.00 | 6 | 0.22 |
| UGT1A1 | 0.03 | 0.00 | 0.00 | 0.00 | 0.05 | 0.00 | 9 | 0.69 |
| UGT2B15 | 0.03 | 0.13 | 0.33 | 0.00 | 0.03 | 0.00 | 15 | 0.62 |
| UGT2B17 | 0.02 | 0.07 | 0.33 | 0.00 | 0.02 | 0.00 | 9 | 0.29 |
| UGT2B7 | 0.06 | 0.12 | 0.33 | 0.00 | 0.04 | 0.00 | 19 | 1.15 |

Table S5: Regions of homozygosity in the extended genes split by group. For each gene the number of ROH found across all samples is shown by group as a fraction of the individuals in that group who share the ROHi, followed the total number and the total normalised by gene length.

| **Gene** | **SA** | **SC** | **KS** | **FW** | **W** | **O** | **# ROHi** | **ROHi/kb** |
|---|---|---|---|---|---|---|---|---|
| ABCA1 | 0.02 | 0.00 | 0.00 | 0.00 | 0.01 | 0.00 | 6 | 0.04 |
| ABCA4 | 0.03 | 0.04 | 0.00 | 0.00 | 0.05 | 0.33 | 18 | 0.14 |
| ABCB11 | 0.02 | 0.04 | 0.00 | 0.00 | 0.01 | 0.00 | 8 | 0.07 |
| ABCB4 | 0.06 | 0.03 | 0.20 | 0.00 | 0.05 | 0.00 | 22 | 0.30 |
| ABCB5 | 0.02 | 0.03 | 0.00 | 0.00 | 0.02 | 0.00 | 9 | 0.06 |
| ABCB6 | 0.02 | 0.01 | 0.20 | 0.00 | 0.01 | 0.00 | 7 | 0.76 |
| ABCB7 | 0.00 | 0.00 | 0.00 | 0.00 | 0.00 | 0.33 | 1 | 0.01 |
| ABCB8 | 0.00 | 0.01 | 0.00 | 0.00 | 0.01 | 0.00 | 3 | 0.15 |
| ABCC1 | 0.01 | 0.00 | 0.00 | 0.00 | 0.00 | 0.00 | 1 | 0.01 |
| ABCC10 | 0.01 | 0.04 | 0.00 | 0.00 | 0.02 | 0.33 | 10 | 0.44 |
| ABCC11 | 0.04 | 0.07 | 0.20 | 0.00 | 0.06 | 0.33 | 26 | 0.38 |
| ABCC12 | 0.07 | 0.12 | 0.20 | 0.00 | 0.10 | 0.67 | 43 | 0.67 |
| ABCC13 | 0.01 | 0.02 | 0.00 | 0.00 | 0.00 | 0.00 | 3 | 0.11 |
| ABCC3 | 0.02 | 0.04 | 0.00 | 0.00 | 0.00 | 0.33 | 8 | 0.14 |
| ABCC4 | 0.03 | 0.01 | 0.00 | 0.00 | 0.01 | 0.00 | 7 | 0.02 |
| ABCC5 | 0.02 | 0.05 | 0.00 | 0.00 | 0.02 | 0.00 | 14 | 0.14 |
| ABCC5-AS1 | 0.02 | 0.05 | 0.00 | 0.00 | 0.02 | 0.00 | 13 | 2.56 |
| ABCC6 | 0.00 | 0.00 | 0.00 | 0.00 | 0.00 | 0.33 | 1 | 0.01 |
| ABCC8 | 0.05 | 0.01 | 0.00 | 0.00 | 0.06 | 0.00 | 18 | 0.21 |
| ABCC9 | 0.06 | 0.05 | 0.00 | 0.00 | 0.04 | 0.00 | 22 | 0.16 |
| ABCG1 | 0.01 | 0.01 | 0.00 | 0.00 | 0.00 | 0.00 | 3 | 0.03 |
| ADH1A | 0.01 | 0.04 | 0.20 | 0.00 | 0.06 | 0.00 | 18 | 1.23 |
| ADH1B | 0.02 | 0.05 | 0.20 | 0.00 | 0.06 | 0.00 | 20 | 1.33 |
| ADH1C | 0.01 | 0.05 | 0.20 | 0.00 | 0.06 | 0.00 | 18 | 1.09 |
| ADH4 | 0.01 | 0.05 | 0.20 | 0.00 | 0.07 | 0.00 | 20 | 0.97 |
| ADH5 | 0.01 | 0.04 | 0.20 | 0.00 | 0.07 | 0.00 | 19 | 1.07 |
| ADH6 | 0.01 | 0.04 | 0.20 | 0.00 | 0.07 | 0.00 | 19 | 1.14 |
| ADH7 | 0.01 | 0.04 | 0.00 | 0.00 | 0.05 | 0.00 | 14 | 0.60 |
| ADHFE1 | 0.02 | 0.02 | 0.00 | 0.14 | 0.03 | 0.33 | 12 | 0.33 |
| AHR | 0.01 | 0.00 | 0.00 | 0.00 | 0.01 | 0.00 | 3 | 0.06 |
| ALDH1A1 | 0.03 | 0.07 | 0.00 | 0.29 | 0.06 | 0.00 | 25 | 0.47 |
| ALDH1A2 | 0.01 | 0.00 | 0.00 | 0.00 | 0.00 | 0.00 | 1 | 0.01 |
| ALDH1A3 | 0.01 | 0.01 | 0.00 | 0.00 | 0.01 | 0.00 | 4 | 0.11 |
| ALDH1B1 | 0.00 | 0.00 | 0.00 | 0.00 | 0.00 | 0.33 | 1 | 0.17 |
| ALDH2 | 0.06 | 0.07 | 0.00 | 0.14 | 0.07 | 0.67 | 32 | 0.74 |
| ALDH3A1 | 0.02 | 0.09 | 0.00 | 0.14 | 0.03 | 0.00 | 19 | 1.82 |
| ALDH3A2 | 0.01 | 0.08 | 0.00 | 0.14 | 0.02 | 0.00 | 16 | 0.55 |
| ALDH3B1 | 0.01 | 0.00 | 0.00 | 0.00 | 0.00 | 0.00 | 2 | 0.10 |
| ALDH3B2 | 0.02 | 0.01 | 0.00 | 0.00 | 0.01 | 0.00 | 6 | 0.31 |
| ALDH4A1 | 0.01 | 0.02 | 0.00 | 0.00 | 0.01 | 0.00 | 5 | 0.16 |
| ALDH5A1 | 0.02 | 0.03 | 0.00 | 0.00 | 0.01 | 0.00 | 7 | 0.17 |

| Gene | SA | SC | KS | FW | W | O | # ROHi | ROHi/kb |
|---|---|---|---|---|---|---|---|---|
| ALDH6A1 | 0.02 | 0.01 | 0.00 | 0.00 | 0.02 | 0.00 | 8 | 0.30 |
| ALDH7A1 | 0.02 | 0.00 | 0.00 | 0.00 | 0.01 | 0.00 | 5 | 0.09 |
| ALDH8A1 | 0.00 | 0.02 | 0.00 | 0.00 | 0.01 | 0.00 | 4 | 0.12 |
| ALDH9A1 | 0.03 | 0.02 | 0.00 | 0.00 | 0.02 | 0.00 | 11 | 0.30 |
| AOX1 | 0.06 | 0.01 | 0.00 | 0.00 | 0.01 | 0.00 | 11 | 0.13 |
| ARNT | 0.05 | 0.07 | 0.20 | 0.00 | 0.04 | 0.00 | 23 | 0.34 |
| ARSA | 0.01 | 0.02 | 0.00 | 0.00 | 0.00 | 0.00 | 4 | 0.74 |
| ATP7A | 0.00 | 0.00 | 0.00 | 0.00 | 0.00 | 0.33 | 1 | 0.01 |
| ATP7B | 0.09 | 0.13 | 0.00 | 0.00 | 0.12 | 0.00 | 48 | 0.61 |
| CAT | 0.01 | 0.01 | 0.00 | 0.00 | 0.01 | 0.00 | 3 | 0.09 |
| CBR1 | 0.01 | 0.00 | 0.00 | 0.00 | 0.02 | 0.00 | 4 | 1.23 |
| CBR3 | 0.01 | 0.01 | 0.00 | 0.00 | 0.01 | 0.00 | 3 | 0.26 |
| CBR3-AS1 | 0.01 | 0.01 | 0.00 | 0.00 | 0.01 | 0.00 | 3 | 0.12 |
| CDA | 0.01 | 0.00 | 0.00 | 0.00 | 0.00 | 0.00 | 2 | 0.07 |
| CES1 | 0.00 | 0.01 | 0.00 | 0.00 | 0.00 | 0.00 | 1 | 0.03 |
| CES2 | 0.07 | 0.11 | 0.00 | 0.00 | 0.10 | 0.33 | 40 | 3.76 |
| CFTR | 0.06 | 0.06 | 0.00 | 0.14 | 0.13 | 0.00 | 39 | 0.21 |
| CHST1 | 0.02 | 0.03 | 0.00 | 0.14 | 0.01 | 0.00 | 9 | 0.50 |
| CHST10 | 0.01 | 0.04 | 0.20 | 0.00 | 0.05 | 0.00 | 15 | 0.58 |
| CHST11 | 0.02 | 0.07 | 0.00 | 0.00 | 0.01 | 0.00 | 13 | 0.04 |
| CHST12 | 0.01 | 0.01 | 0.00 | 0.00 | 0.00 | 0.00 | 3 | 0.10 |
| CHST13 | 0.01 | 0.02 | 0.20 | 0.00 | 0.02 | 0.33 | 10 | 0.53 |
| CHST2 | 0.01 | 0.01 | 0.00 | 0.00 | 0.00 | 0.00 | 3 | 0.71 |
| CHST3 | 0.02 | 0.04 | 0.00 | 0.00 | 0.02 | 0.00 | 10 | 0.20 |
| CHST4 | 0.03 | 0.07 | 0.20 | 0.14 | 0.10 | 0.00 | 31 | 2.49 |
| CHST5 | 0.02 | 0.06 | 0.20 | 0.14 | 0.04 | 0.00 | 19 | 2.86 |
| CHST6 | 0.02 | 0.08 | 0.20 | 0.14 | 0.03 | 0.00 | 20 | 0.91 |
| CHST7 | 0.00 | 0.00 | 0.00 | 0.00 | 0.00 | 0.33 | 1 | 0.04 |
| CHST8 | 0.02 | 0.02 | 0.00 | 0.00 | 0.00 | 0.00 | 5 | 0.03 |
| CHST9 | 0.02 | 0.04 | 0.20 | 0.00 | 0.01 | 0.00 | 9 | 0.03 |
| CYB5R3 | 0.01 | 0.00 | 0.00 | 0.00 | 0.00 | 0.00 | 2 | 0.06 |
| CYP11A1 | 0.03 | 0.03 | 0.00 | 0.00 | 0.05 | 0.00 | 16 | 0.53 |
| CYP11B1 | 0.01 | 0.00 | 0.00 | 0.00 | 0.01 | 0.00 | 2 | 0.27 |
| CYP11B2 | 0.01 | 0.00 | 0.00 | 0.00 | 0.01 | 0.00 | 2 | 0.27 |
| CYP17A1 | 0.18 | 0.27 | 0.20 | 0.00 | 0.27 | 0.33 | 105 | 14.99 |
| CYP19A1 | 0.03 | 0.04 | 0.00 | 0.00 | 0.07 | 0.00 | 22 | 0.17 |
| CYP1B1 | 0.02 | 0.00 | 0.20 | 0.00 | 0.02 | 0.00 | 8 | 0.93 |
| CYP1B1-AS1 | 0.01 | 0.00 | 0.20 | 0.00 | 0.04 | 0.00 | 9 | 0.18 |
| CYP20A1 | 0.11 | 0.09 | 0.00 | 0.00 | 0.11 | 0.67 | 47 | 0.70 |
| CYP21A2 | 0.07 | 0.06 | 0.00 | 0.14 | 0.02 | 0.00 | 23 | 6.86 |
| CYP24A1 | 0.01 | 0.00 | 0.00 | 0.00 | 0.01 | 0.00 | 2 | 0.10 |
| CYP26A1 | 0.02 | 0.03 | 0.00 | 0.00 | 0.02 | 0.00 | 10 | 2.27 |
| CYP26C1 | 0.02 | 0.04 | 0.00 | 0.00 | 0.02 | 0.00 | 11 | 1.48 |
| CYP27B1 | 0.04 | 0.06 | 0.00 | 0.00 | 0.02 | 0.00 | 17 | 3.50 |
| CYP2A13 | 0.00 | 0.00 | 0.00 | 0.00 | 0.00 | 0.33 | 1 | 0.13 |

| Gene | SA | SC | KS | FW | W | O | # ROHi | ROHi/kb |
|---|---|---|---|---|---|---|---|---|
| CYP2A7 | 0.00 | 0.00 | 0.00 | 0.00 | 0.00 | 0.33 | 1 | 0.14 |
| CYP2C18 | 0.11 | 0.08 | 0.00 | 0.14 | 0.12 | 0.00 | 46 | 0.87 |
| CYP2F1 | 0.00 | 0.00 | 0.00 | 0.00 | 0.00 | 0.33 | 1 | 0.07 |
| CYP2J2 | 0.03 | 0.04 | 0.00 | 0.00 | 0.01 | 0.00 | 11 | 0.33 |
| CYP2R1 | 0.11 | 0.10 | 0.00 | 0.14 | 0.09 | 0.00 | 43 | 3.03 |
| CYP2S1 | 0.01 | 0.00 | 0.00 | 0.00 | 0.00 | 0.00 | 1 | 0.07 |
| CYP39A1 | 0.03 | 0.04 | 0.00 | 0.00 | 0.03 | 0.33 | 15 | 0.15 |
| CYP3A43 | 0.07 | 0.04 | 0.00 | 0.00 | 0.03 | 0.00 | 21 | 0.55 |
| CYP3A7 | 0.06 | 0.05 | 0.20 | 0.14 | 0.06 | 0.00 | 27 | 0.90 |
| CYP3A7-CYP3AP1 | 0.06 | 0.05 | 0.20 | 0.14 | 0.06 | 0.00 | 27 | 0.53 |
| CYP46A1 | 0.02 | 0.01 | 0.00 | 0.00 | 0.00 | 0.00 | 4 | 0.09 |
| CYP4A11 | 0.00 | 0.00 | 0.00 | 0.00 | 0.00 | 0.33 | 1 | 0.08 |
| CYP4B1 | 0.00 | 0.00 | 0.00 | 0.00 | 0.00 | 0.33 | 1 | 0.05 |
| CYP4F11 | 0.01 | 0.00 | 0.00 | 0.14 | 0.01 | 0.00 | 4 | 0.18 |
| CYP4F12 | 0.01 | 0.00 | 0.00 | 0.14 | 0.01 | 0.00 | 5 | 0.21 |
| CYP4F2 | 0.01 | 0.00 | 0.00 | 0.14 | 0.01 | 0.00 | 4 | 0.20 |
| CYP4F3 | 0.01 | 0.00 | 0.00 | 0.14 | 0.01 | 0.00 | 5 | 0.25 |
| CYP4F8 | 0.01 | 0.00 | 0.00 | 0.00 | 0.01 | 0.00 | 4 | 0.28 |
| CYP4Z1 | 0.01 | 0.00 | 0.00 | 0.00 | 0.00 | 0.00 | 2 | 0.04 |
| CYP51A1 | 0.09 | 0.09 | 0.00 | 0.29 | 0.09 | 0.33 | 42 | 1.86 |
| CYP7A1 | 0.01 | 0.01 | 0.00 | 0.00 | 0.02 | 0.00 | 7 | 0.70 |
| CYP7B1 | 0.02 | 0.06 | 0.00 | 0.00 | 0.07 | 0.00 | 22 | 0.11 |
| CYP8B1 | 0.01 | 0.06 | 0.00 | 0.00 | 0.04 | 0.00 | 15 | 3.80 |
| DDO | 0.01 | 0.03 | 0.00 | 0.00 | 0.01 | 0.00 | 7 | 0.30 |
| DHRS1 | 0.01 | 0.02 | 0.00 | 0.00 | 0.01 | 0.00 | 4 | 0.43 |
| DHRS12 | 0.04 | 0.06 | 0.00 | 0.00 | 0.05 | 0.00 | 21 | 0.58 |
| DHRS13 | 0.05 | 0.09 | 0.00 | 0.00 | 0.03 | 0.00 | 23 | 4.35 |
| DHRS2 | 0.01 | 0.02 | 0.00 | 0.00 | 0.00 | 0.00 | 4 | 0.43 |
| DHRS3 | 0.01 | 0.02 | 0.20 | 0.00 | 0.02 | 0.00 | 9 | 0.18 |
| DHRS4 | 0.00 | 0.00 | 0.00 | 0.00 | 0.00 | 0.33 | 1 | 0.06 |
| DHRS4-AS1 | 0.00 | 0.00 | 0.00 | 0.00 | 0.00 | 0.33 | 1 | 0.06 |
| DHRS4L1 | 0.01 | 0.02 | 0.00 | 0.00 | 0.01 | 0.00 | 4 | 0.09 |
| DHRS4L2 | 0.01 | 0.02 | 0.00 | 0.00 | 0.01 | 0.00 | 4 | 0.11 |
| DHRS7 | 0.08 | 0.12 | 0.20 | 0.14 | 0.11 | 0.00 | 46 | 2.22 |
| DHRS7B | 0.01 | 0.03 | 0.00 | 0.00 | 0.00 | 0.00 | 4 | 0.06 |
| DHRS7C | 0.02 | 0.00 | 0.00 | 0.00 | 0.00 | 0.00 | 4 | 0.20 |
| DHRS9 | 0.01 | 0.03 | 0.00 | 0.00 | 0.01 | 0.00 | 6 | 0.21 |
| DHRSX | 0.00 | 0.00 | 0.00 | 0.00 | 0.00 | 1.00 | 3 | 0.01 |
| DPEP1 | 0.01 | 0.02 | 0.20 | 0.00 | 0.01 | 0.00 | 6 | 0.24 |
| EPHX1 | 0.02 | 0.01 | 0.20 | 0.00 | 0.02 | 0.00 | 8 | 0.23 |
| EPHX2 | 0.03 | 0.00 | 0.00 | 0.00 | 0.01 | 0.00 | 7 | 0.13 |
| FMO1 | 0.02 | 0.00 | 0.00 | 0.00 | 0.02 | 0.00 | 6 | 0.16 |
| FMO2 | 0.02 | 0.00 | 0.00 | 0.00 | 0.02 | 0.00 | 6 | 0.22 |
| FMO3 | 0.02 | 0.00 | 0.00 | 0.00 | 0.02 | 0.00 | 6 | 0.22 |
| FMO4 | 0.01 | 0.00 | 0.00 | 0.00 | 0.01 | 0.00 | 4 | 0.14 |

| Gene | SA | SC | KS | FW | W | O | # ROHi | ROHi/kb |
|---|---|---|---|---|---|---|---|---|
| FMO5 | 0.04 | 0.02 | 0.00 | 0.00 | 0.04 | 0.00 | 14 | 0.34 |
| FMO6P | 0.02 | 0.00 | 0.00 | 0.00 | 0.02 | 0.00 | 6 | 0.25 |
| GPX1 | 0.05 | 0.10 | 0.40 | 0.14 | 0.05 | 0.33 | 31 | 26.20 |
| GPX2 | 0.04 | 0.02 | 0.00 | 0.00 | 0.02 | 0.00 | 13 | 3.46 |
| GPX3 | 0.01 | 0.00 | 0.00 | 0.14 | 0.01 | 0.00 | 3 | 0.35 |
| GPX4 | 0.00 | 0.00 | 0.20 | 0.00 | 0.00 | 0.00 | 1 | 0.35 |
| GPX5 | 0.04 | 0.10 | 0.20 | 0.29 | 0.09 | 0.00 | 35 | 3.91 |
| GPX6 | 0.04 | 0.10 | 0.20 | 0.29 | 0.09 | 0.00 | 36 | 2.88 |
| GPX7 | 0.04 | 0.04 | 0.20 | 0.00 | 0.01 | 0.00 | 12 | 1.80 |
| GSR | 0.02 | 0.01 | 0.20 | 0.00 | 0.02 | 0.33 | 9 | 0.18 |
| GSS | 0.11 | 0.11 | 0.00 | 0.00 | 0.10 | 0.00 | 46 | 1.68 |
| GSTA1 | 0.01 | 0.00 | 0.00 | 0.00 | 0.00 | 0.00 | 2 | 0.16 |
| GSTA2 | 0.01 | 0.01 | 0.00 | 0.00 | 0.00 | 0.00 | 2 | 0.15 |
| GSTA3 | 0.01 | 0.01 | 0.00 | 0.00 | 0.00 | 0.33 | 4 | 0.31 |
| GSTA4 | 0.02 | 0.04 | 0.00 | 0.14 | 0.01 | 0.33 | 11 | 0.63 |
| GSTA5 | 0.01 | 0.01 | 0.00 | 0.00 | 0.00 | 0.33 | 4 | 0.28 |
| GSTCD | 0.02 | 0.06 | 0.00 | 0.14 | 0.07 | 0.33 | 23 | 0.17 |
| GSTK1 | 0.04 | 0.05 | 0.00 | 0.00 | 0.04 | 0.00 | 20 | 3.51 |
| GSTM2 | 0.00 | 0.03 | 0.00 | 0.00 | 0.01 | 0.00 | 4 | 0.25 |
| GSTM3 | 0.01 | 0.02 | 0.00 | 0.00 | 0.00 | 0.00 | 3 | 0.42 |
| GSTM4 | 0.00 | 0.03 | 0.00 | 0.00 | 0.01 | 0.00 | 5 | 0.53 |
| GSTM5 | 0.00 | 0.02 | 0.00 | 0.00 | 0.01 | 0.00 | 3 | 0.50 |
| GSTO1 | 0.01 | 0.00 | 0.00 | 0.00 | 0.01 | 0.00 | 3 | 0.23 |
| GSTO2 | 0.01 | 0.00 | 0.00 | 0.00 | 0.02 | 0.00 | 4 | 0.13 |
| GSTT2 | 0.01 | 0.04 | 0.00 | 0.00 | 0.02 | 0.00 | 9 | 2.37 |
| GSTZ1 | 0.02 | 0.01 | 0.00 | 0.14 | 0.01 | 0.00 | 8 | 0.75 |
| HAGH | 0.02 | 0.02 | 0.00 | 0.00 | 0.00 | 0.00 | 6 | 0.33 |
| HNF4A | 0.02 | 0.04 | 0.00 | 0.00 | 0.01 | 0.00 | 10 | 0.13 |
| HNF4A-AS1 | 0.02 | 0.03 | 0.00 | 0.00 | 0.01 | 0.00 | 7 | 0.39 |
| HNMT | 0.01 | 0.06 | 0.00 | 0.00 | 0.01 | 0.00 | 10 | 0.19 |
| HSD11B1 | 0.01 | 0.01 | 0.20 | 0.00 | 0.01 | 0.00 | 6 | 0.12 |
| HSD17B11 | 0.04 | 0.03 | 0.00 | 0.00 | 0.01 | 0.00 | 12 | 0.22 |
| HSD17B14 | 0.00 | 0.01 | 0.20 | 0.00 | 0.01 | 0.00 | 3 | 0.13 |
| IAPP | 0.02 | 0.02 | 0.00 | 0.00 | 0.01 | 0.00 | 8 | 1.12 |
| KCNJ11 | 0.05 | 0.02 | 0.00 | 0.00 | 0.06 | 0.00 | 19 | 4.65 |
| MAT1A | 0.02 | 0.02 | 0.00 | 0.00 | 0.02 | 0.00 | 9 | 0.50 |
| METAP1 | 0.01 | 0.04 | 0.20 | 0.00 | 0.06 | 0.00 | 17 | 0.25 |
| MGST1 | 0.02 | 0.00 | 0.00 | 0.00 | 0.01 | 0.00 | 4 | 0.13 |
| MGST2 | 0.01 | 0.01 | 0.20 | 0.00 | 0.01 | 0.00 | 4 | 0.05 |
| MGST3 | 0.03 | 0.02 | 0.00 | 0.00 | 0.02 | 0.00 | 11 | 0.44 |
| MPO | 0.02 | 0.03 | 0.00 | 0.00 | 0.01 | 0.00 | 8 | 0.72 |
| NNMT | 0.01 | 0.01 | 0.00 | 0.00 | 0.01 | 0.00 | 4 | 0.24 |
| NOS1 | 0.01 | 0.01 | 0.00 | 0.00 | 0.01 | 0.00 | 3 | 0.02 |
| NOS2 | 0.01 | 0.03 | 0.00 | 0.00 | 0.03 | 0.00 | 9 | 0.21 |
| NOS3 | 0.00 | 0.01 | 0.00 | 0.00 | 0.01 | 0.00 | 3 | 0.13 |

| Gene | SA | SC | KS | FW | W | O | # ROHi | ROHi/kb |
|---|---|---|---|---|---|---|---|---|
| NR1I2 | 0.07 | 0.09 | 0.00 | 0.00 | 0.14 | 0.00 | 45 | 1.18 |
| NR1I3 | 0.01 | 0.01 | 0.00 | 0.00 | 0.04 | 0.00 | 10 | 1.17 |
| PDE3A | 0.02 | 0.03 | 0.20 | 0.00 | 0.02 | 0.00 | 10 | 0.03 |
| PDE3B | 0.14 | 0.11 | 0.00 | 0.14 | 0.12 | 0.00 | 55 | 0.24 |
| PLGLB1 | 0.00 | 0.00 | 0.00 | 0.00 | 0.00 | 0.67 | 2 | 0.18 |
| PNMT | 0.08 | 0.06 | 0.00 | 0.57 | 0.06 | 0.00 | 34 | 13.63 |
| PON1 | 0.07 | 0.02 | 0.00 | 0.00 | 0.02 | 0.00 | 17 | 0.65 |
| PON2 | 0.07 | 0.01 | 0.00 | 0.00 | 0.01 | 0.00 | 14 | 0.46 |
| PON3 | 0.07 | 0.01 | 0.00 | 0.00 | 0.02 | 0.00 | 15 | 0.41 |
| POR | 0.01 | 0.02 | 0.00 | 0.00 | 0.01 | 0.00 | 4 | 0.06 |
| PPARA | 0.02 | 0.02 | 0.00 | 0.00 | 0.00 | 0.00 | 5 | 0.05 |
| PPARD | 0.06 | 0.05 | 0.20 | 0.00 | 0.07 | 0.00 | 27 | 0.32 |
| PPARG | 0.06 | 0.08 | 0.20 | 0.00 | 0.06 | 0.00 | 28 | 0.19 |
| RXRA | 0.00 | 0.00 | 0.00 | 0.00 | 0.00 | 0.33 | 1 | 0.01 |
| SERPINA7 | 0.00 | 0.00 | 0.00 | 0.00 | 0.00 | 0.33 | 1 | 0.18 |
| SLC10A1 | 0.02 | 0.01 | 0.00 | 0.00 | 0.02 | 0.00 | 7 | 0.33 |
| SLC10A2 | 0.00 | 0.00 | 0.00 | 0.00 | 0.01 | 0.00 | 2 | 0.09 |
| SLC13A1 | 0.06 | 0.05 | 0.20 | 0.00 | 0.06 | 0.00 | 26 | 0.30 |
| SLC13A2 | 0.02 | 0.03 | 0.00 | 0.00 | 0.04 | 0.00 | 13 | 0.54 |
| SLC13A3 | 0.01 | 0.00 | 0.20 | 0.00 | 0.00 | 0.00 | 2 | 0.02 |
| SLC15A1 | 0.01 | 0.01 | 0.00 | 0.00 | 0.01 | 0.00 | 4 | 0.06 |
| SLC16A1 | 0.02 | 0.05 | 0.00 | 0.00 | 0.03 | 0.00 | 14 | 0.31 |
| SLC16A1-AS1 | 0.02 | 0.05 | 0.00 | 0.00 | 0.03 | 0.00 | 14 | 1.83 |
| SLC19A1 | 0.01 | 0.01 | 0.00 | 0.00 | 0.01 | 0.00 | 5 | 0.18 |
| SLC22A10 | 0.04 | 0.02 | 0.00 | 0.00 | 0.03 | 0.33 | 14 | 0.64 |
| SLC22A11 | 0.04 | 0.01 | 0.00 | 0.00 | 0.02 | 0.00 | 11 | 0.69 |
| SLC22A12 | 0.04 | 0.00 | 0.00 | 0.00 | 0.02 | 0.00 | 10 | 0.87 |
| SLC22A13 | 0.04 | 0.04 | 0.00 | 0.00 | 0.06 | 0.00 | 21 | 1.68 |
| SLC22A14 | 0.04 | 0.04 | 0.00 | 0.00 | 0.04 | 0.00 | 18 | 1.45 |
| SLC22A15 | 0.04 | 0.05 | 0.20 | 0.00 | 0.04 | 0.33 | 21 | 0.22 |
| SLC22A16 | 0.02 | 0.03 | 0.00 | 0.00 | 0.02 | 0.00 | 9 | 0.17 |
| SLC22A17 | 0.02 | 0.01 | 0.00 | 0.00 | 0.00 | 0.00 | 5 | 0.76 |
| SLC22A18 | 0.01 | 0.01 | 0.00 | 0.00 | 0.02 | 0.00 | 6 | 0.24 |
| SLC22A18AS | 0.01 | 0.01 | 0.00 | 0.00 | 0.02 | 0.00 | 6 | 0.38 |
| SLC22A3 | 0.00 | 0.01 | 0.00 | 0.00 | 0.03 | 0.00 | 6 | 0.06 |
| SLC22A4 | 0.04 | 0.00 | 0.00 | 0.00 | 0.01 | 0.00 | 8 | 0.16 |
| SLC22A5 | 0.04 | 0.01 | 0.00 | 0.00 | 0.01 | 0.00 | 9 | 0.35 |
| SLC22A7 | 0.02 | 0.04 | 0.00 | 0.00 | 0.02 | 0.33 | 12 | 1.65 |
| SLC22A8 | 0.04 | 0.01 | 0.00 | 0.00 | 0.03 | 0.00 | 13 | 0.56 |
| SLC22A9 | 0.03 | 0.01 | 0.00 | 0.00 | 0.03 | 0.33 | 12 | 0.30 |
| SLC27A1 | 0.01 | 0.01 | 0.00 | 0.00 | 0.00 | 0.00 | 3 | 0.08 |
| SLC28A1 | 0.02 | 0.01 | 0.20 | 0.00 | 0.02 | 0.00 | 9 | 0.15 |
| SLC28A2 | 0.01 | 0.04 | 0.00 | 0.00 | 0.02 | 0.00 | 10 | 0.42 |
| SLC28A3 | 0.01 | 0.01 | 0.40 | 0.00 | 0.01 | 0.00 | 7 | 0.08 |
| SLC29A1 | 0.02 | 0.01 | 0.00 | 0.00 | 0.00 | 0.33 | 5 | 0.34 |

| Gene | SA | SC | KS | FW | W | O | # ROHi | ROHi/kb |
|---|---|---|---|---|---|---|---|---|
| SLC29A2 | 0.04 | 0.04 | 0.00 | 0.14 | 0.02 | 0.33 | 17 | 1.83 |
| SLC2A4 | 0.01 | 0.00 | 0.00 | 0.00 | 0.00 | 0.00 | 1 | 0.16 |
| SLC2A5 | 0.01 | 0.01 | 0.00 | 0.00 | 0.01 | 0.00 | 3 | 0.09 |
| SLC5A6 | 0.04 | 0.03 | 0.40 | 0.00 | 0.05 | 0.33 | 20 | 1.57 |
| SLC6A6 | 0.00 | 0.02 | 0.00 | 0.00 | 0.00 | 0.00 | 2 | 0.02 |
| SLC7A5 | 0.01 | 0.00 | 0.20 | 0.00 | 0.00 | 0.00 | 2 | 0.05 |
| SLC7A7 | 0.01 | 0.01 | 0.00 | 0.00 | 0.00 | 0.00 | 3 | 0.06 |
| SLC7A8 | 0.02 | 0.02 | 0.00 | 0.00 | 0.00 | 0.00 | 6 | 0.10 |
| SLCO1A2 | 0.03 | 0.02 | 0.00 | 0.00 | 0.02 | 0.00 | 11 | 0.08 |
| SLCO1C1 | 0.03 | 0.02 | 0.20 | 0.00 | 0.02 | 0.00 | 12 | 0.21 |
| SLCO2A1 | 0.03 | 0.03 | 0.20 | 0.00 | 0.02 | 0.33 | 13 | 0.13 |
| SLCO2B1 | 0.04 | 0.03 | 0.00 | 0.00 | 0.01 | 0.00 | 10 | 0.18 |
| SLCO3A1 | 0.01 | 0.03 | 0.00 | 0.00 | 0.02 | 0.00 | 8 | 0.03 |
| SLCO4A1 | 0.01 | 0.00 | 0.00 | 0.00 | 0.01 | 0.00 | 4 | 0.13 |
| SLCO4C1 | 0.04 | 0.10 | 0.00 | 0.29 | 0.09 | 0.33 | 36 | 0.58 |
| SLCO5A1 | 0.01 | 0.01 | 0.00 | 0.00 | 0.01 | 0.00 | 4 | 0.02 |
| SLCO6A1 | 0.03 | 0.07 | 0.00 | 0.29 | 0.08 | 0.33 | 29 | 0.23 |
| SLX1A-SULT1A3 | 0.01 | 0.04 | 0.00 | 0.00 | 0.01 | 0.33 | 8 | 0.81 |
| SOD1 | 0.03 | 0.03 | 0.00 | 0.00 | 0.01 | 0.00 | 9 | 0.97 |
| SOD2 | 0.02 | 0.02 | 0.00 | 0.00 | 0.01 | 0.00 | 6 | 0.42 |
| SOD3 | 0.01 | 0.00 | 0.20 | 0.00 | 0.01 | 0.33 | 5 | 0.93 |
| SULF1 | 0.02 | 0.00 | 0.00 | 0.00 | 0.01 | 0.00 | 5 | 0.03 |
| SULT1A2 | 0.00 | 0.00 | 0.00 | 0.00 | 0.00 | 0.33 | 1 | 0.20 |
| SULT1A3 | 0.01 | 0.04 | 0.00 | 0.00 | 0.01 | 0.33 | 8 | 1.57 |
| SULT1B1 | 0.01 | 0.01 | 0.20 | 0.00 | 0.06 | 0.00 | 13 | 0.39 |
| SULT1C2 | 0.04 | 0.01 | 0.20 | 0.14 | 0.03 | 0.00 | 14 | 0.66 |
| SULT1E1 | 0.01 | 0.02 | 0.20 | 0.00 | 0.08 | 0.00 | 18 | 0.95 |
| SULT2A1 | 0.00 | 0.02 | 0.20 | 0.00 | 0.00 | 0.00 | 3 | 0.19 |
| SULT2B1 | 0.01 | 0.02 | 0.20 | 0.00 | 0.01 | 0.00 | 6 | 0.13 |
| SULT4A1 | 0.02 | 0.04 | 0.20 | 0.14 | 0.06 | 0.00 | 20 | 0.53 |
| TAP1 | 0.02 | 0.02 | 0.00 | 0.00 | 0.01 | 0.00 | 6 | 0.68 |
| TAP2 | 0.02 | 0.03 | 0.00 | 0.00 | 0.01 | 0.00 | 8 | 0.47 |
| UGT1A10 | 0.02 | 0.01 | 0.00 | 0.00 | 0.04 | 0.00 | 10 | 0.07 |
| UGT1A3 | 0.02 | 0.00 | 0.00 | 0.00 | 0.04 | 0.00 | 9 | 0.20 |
| UGT1A4 | 0.02 | 0.01 | 0.00 | 0.00 | 0.04 | 0.00 | 10 | 0.18 |
| UGT1A5 | 0.02 | 0.01 | 0.00 | 0.00 | 0.04 | 0.00 | 10 | 0.17 |
| UGT1A6 | 0.02 | 0.01 | 0.00 | 0.00 | 0.04 | 0.00 | 10 | 0.12 |
| UGT1A7 | 0.02 | 0.01 | 0.00 | 0.00 | 0.04 | 0.00 | 10 | 0.11 |
| UGT1A8 | 0.02 | 0.01 | 0.00 | 0.00 | 0.04 | 0.00 | 10 | 0.06 |
| UGT1A9 | 0.02 | 0.01 | 0.00 | 0.00 | 0.04 | 0.00 | 10 | 0.10 |
| UGT2A1 | 0.01 | 0.03 | 0.20 | 0.00 | 0.05 | 0.00 | 14 | 0.22 |
| UGT2B10 | 0.04 | 0.17 | 0.40 | 0.00 | 0.10 | 0.00 | 45 | 2.81 |
| UGT2B11 | 0.03 | 0.04 | 0.20 | 0.00 | 0.01 | 0.00 | 13 | 0.90 |
| UGT2B28 | 0.04 | 0.01 | 0.20 | 0.00 | 0.02 | 0.00 | 11 | 0.76 |
| UGT2B4 | 0.01 | 0.03 | 0.20 | 0.00 | 0.02 | 0.00 | 10 | 0.64 |

| Gene | SA | SC | KS | FW | W | O | # ROHi | ROHi/kb |
|---|---|---|---|---|---|---|---|---|
| UGT8 | 0.05 | 0.04 | 0.00 | 0.00 | 0.06 | 0.00 | 22 | 0.28 |
| XDH | 0.06 | 0.05 | 0.20 | 0.00 | 0.06 | 0.33 | 27 | 0.34 |

#### References


[1] Choudhury A, Aron S, Botigué L, Sengupta D, Botha G, Bensellak T, Wells G, Kumuthini J, Shriner D, Fakim YJ, Ghoorah AW, Dareng E, Odia T, Falola O, Adebiyi E, Hazelhurst S, Mazandu G, Nyangiri OA, Mbiyavanga M, Benkahla A, Kassim SK, Mulder N, Adebamowo SN, Chimusa ER, Gibbs RA, Muzny D, Metcalf G, TrypanoGEN Research Group, Rotimi C, Ramsay M, Adeyemo A, Lombard Z, Hanchard NA as members of the H3Africa Consortium. High Depth African Genomes Inform Human Migration and Health. In Preparation

[2] Alexander DH, Novembre J, Lange K. Fast model-based estimation of ancestry in unrelated individuals. Genome Research. 2009;19(9):1655–64. doi:10.1101/gr.094052.109.

[3] Jakobsson M, Rosenberg NA. CLUMPP: a cluster matching and permutation program for dealing with label switching and multimodality in analysis of population structure. Bioinformatics. 2007;23(14):1801–1806.

[4] Rentzsch P, Witten D, Cooper GM, Shendure J, Kircher M. CADD: Predicting the deleteriousness of variants throughout the human genome. Nucleic Acids Research. 2019;47(D1):D886–D894. doi:10.1093/nar/gky1016.
